# Spatial Transcriptomics Analysis Uncovers ER stress in MANF-deficient Purkinje Cells Underlying Alcohol-induced Cerebellar Vulnerability in Mice

**DOI:** 10.1101/2025.06.19.660571

**Authors:** Wen Wen, Hui Li, Li-Chun Lin, Michael S. Chimenti, Henry L. Keen, Mariah R. Leidinger, Di Hu, Zuohui Zhang, Hong Lin, Jia Luo

**Affiliations:** Department of Pathology, Carver College of Medicine, University of Iowa, Iowa City, IA 52242, USA; Iowa Neuroscience Institute, Carver College of Medicine, University of Iowa, Iowa City, IA 52242, USA; Bioinformatics Division, Iowa Institute of Human Genetics, Carver College of Medicine, University of Iowa, Iowa City, IA 52242, USA; Comparative Pathology Laboratory, Carver College of Medicine, University of Iowa, Iowa City, IA 52242, USA; Iowa City VA Health Care System, Iowa City, IA 52246, USA

**Keywords:** Alcohol use disorder, ER stress, unfolded protein response, neurodegeneration, sex difference, transcriptomics

## Abstract

Cerebellar Purkinje cells (PCs) are among the most vulnerable neurons to alcohol neurotoxicity. Alcohol can induce endoplasmic reticulum (ER) stress and alter the structure and function of PCs. Mesencephalic astrocyte-derived neurotrophic factor (MANF) is an ER stress inducible protein highly expressed in PCs. It is neuroprotective in various pathological conditions where ER stress is induced. However, it is unknown whether MANF plays a role in protecting PCs from alcohol induced ER stress. In this study, we generated PC-specific MANF knockout (KO) mouse model to test the hypothesis that MANF deficient PCs are more susceptible to binge alcohol exposure induced ER stress and neurodegeneration in the adult brain. We found that PC-specific MANF KO animals show moderate motor function deficit, which was exacerbated by alcohol exposure. Interestingly, female KOs were more sensitive than male KOs to alcohol-induced motor function impairments. In accordance with the behavior changes, alcohol exposure also caused UPR activation, increased intranuclear expression of calcium binding protein Calbindin, and PC degeneration in female but not male MANF KO mice. Spatial transcriptomics and high throughput in situ analyses demonstrated that MANF deficiency altered the transcriptomic landscape in PCs in a sex-specific manner and triggered the expression of genes involved in protein folding and response to ER stress. These results suggests that MANF KO PCs may be predisposed with a higher risk to UPR activation and ER stress in a sex dependent manner, contributing to their vulnerability to alcohol neurotoxicity.

## Introduction

Excessive alcohol consumption has profound adverse effects on the brain, affecting both neuronal structure and function. The cerebellum is one of the most vulnerable brain regions to the toxic effects of alcohol [1]. Even low levels of acute alcohol intoxication can impair cerebellar function, leading to noticeable motor deficits such as unsteady gait and poor coordination [2]. Chronic alcohol consumption and binge drinking can lead to irreversible cerebellar degeneration, which is one of the most common neurological complications in patients with alcohol use disorder. The cerebellum is not only responsible for coordinating motor movements, but also involves in non-motor functions such as memory, language, and emotional control. As a result, alcohol induced cerebellar degeneration can lead to long-term motor impairments and may also play a significant role in the adverse impact of alcohol on cognition and emotion [3]. Excessive alcohol exposure is associated with structural abnormalities and neurologic cerebellar dysfunction with anterior and superior vermis atrophy, decrease in the volume of molecular and granular layers, and loss of Purkinje cells (PCs) [4, 5]. PCs are the sole efferent neurons of the cerebellum, sending output from the cerebellar cortex to the cerebral cortex and brain stem. Alcohol exposure during early development or chronic alcohol exposure in adult can alter the structure and function of PCs [6–13]. Reduced numbers of PCs are often reported in postmortem human brains in individuals with chronic alcohol consumption [14–20]. Animal studies indicate that alcohol exposure can cause PC shrinkage, dendritic arbor reduction, mitochondrial fragmentation, disruption in Ca^2+^ signaling, synaptic impairment, electrophysiological changes, and neurotransmission alteration [6, 7, 13, 21–24].

The mechanisms by which alcohol causes cerebellar degeneration are complex and multifaceted, involving direct neurotoxic effects of ethanol and its metabolic products, nutritional deficiencies, such as thiamine deficiency, potential immune-mediated processes, and dysregulation of neurotrophic factors [5]. Understanding the impact of alcohol on the cerebellum is crucial for developing effective interventions and treatments for alcohol-related brain damages. Alcohol induces endoplasmic reticulum (ER) stress which is associated with multiple organ damages [25]. We and other investigators have demonstrated that alcohol exposure induces ER stress in both the developing and mature brain, including the cerebellum [8, 26–35]. The ER is an important organelle for the biological function and homeostasis of cells, regulating posttranslational modification, folding, and transportation of proteins, and the synthesis of membrane lipids and cholesterol. ER chaperone molecules such as the 78-kDa glucose-regulated protein (GRP78) facilitates protein folding to maintain ER protein homeostasis in non-stressed condition. Under stress conditions such as pathogen infection, nutrient deprivation, and inflammation, the abundance of misfolded proteins access the capacity of the protein folding machinery in the ER, the unfolded protein response (UPR) will be triggered via the activation of three ER transmembrane receptors: pancreatic ER kinase-like ER kinase (PERK), inositol-requiring enzyme 1 (IRE1), and activating transcription factor 6 (ATF6) [36]. UPR alleviates ER-stress by suppressing the rate of protein synthesis, degrading the unfolded or misfolded proteins through ER-associated protein degradation (ERAD), and producing molecular chaperons to increase the protein folding capacity in the ER [36]. If the UPR is insufficient to overcome ER-stress, prolonged ER-stress can ultimately results in apoptosis [37]. Chronic alcohol exposure has been reported to increase the expression of ER stress markers in the PCs and induce dilation of the smooth endoplasmic reticulum (SER) which precedes and accompanies dendritic regression of PCs in rats [7–9].

Mesencephalic astrocyte-derived neurotrophic factor (MANF) is an evolutionarily conserved ER resident protein belonging to a novel neurotrophic factor family [38]. It was originally identified as a secreted trophic factor for dopamine neurons in vitro [39]. It is systemically expressed throughout the nerves and non-nervous systems. In the rodent brain, robust MANF expression can be detected in neurons from embryonic stages to adulthood and declined during aging, suggesting its role in neurodevelopment and maintenance of neuronal functions [40–42]. MANF has a cytoprotective role in regulating UPR and maintaining ER homeostasis [43, 44]. MANF protein physically interacts with the major ER chaperone GRP78 [45]. Constitutive MANF-deficient mouse exhibit chronic UPR activation in the brain [46]. MANF expression and secretion are upregulated in response to ER stress [47–49]. Generally, MANF upregulation offers protection from ER stress in experimental models of Parkinson’s disease, brain and heart ischemia, and retinal degeneration [41, 50–55].

Previously, we have demonstrated that acute alcohol exposure caused significant upregulation of MANF in the developing mice brain which was accompanied with increased ER-stress [26, 33]. move this sentence to discussion. MANF is highly expressed in PCs in the adult mouse brain [41]. It has been reported to protect and restore damaged PCs in mouse model of spinocerebellar ataxia (SCA) [56]. However, whether MANF protects PCs from alcohol-induced degeneration is unknown. In the present study, we generated a PC-specific MANF KO mouse model to test the hypothesis that MANF deficient PCs are more susceptible to alcohol induced ER stress and neurodegeneration. Furthermore, we examined the transcriptomic changes due to MANF deficiency in PCs to identify genes and signaling pathways that potentially interacted with MANF using Visium spatial transcriptomics analysis (STA) followed by Xenium in situ analysis.

## Materials and Methods

### Animal husbandry

All animals were group housed (up to 5 per cage) and cared for by the Office of Animal Resources at the University of Iowa. Mice were allowed ad libitum access to chow and water on a 12-h light/12-h dark cycle. All experimental animal procedures were approved by the Institutional Animal Care and Use Committee (IACUC) at the University of Iowa (#3042295) and performed following regulations for the Care and Use of Laboratory Animals set forth by the National Institutes of Health (NIH) Guide.

### Mouse strains and genotyping

Adult male and female mice were used for this study. *Manf* ^fl^/^fl^ transgenic mice with C57BL/6 background were generated as described previously [31, 32]. Purkinje cell specific Cre mice B6.Cg-Tg(Pcp2-cre)3555Jdhu/J (referred to as *Pcp2-Cre* and thereafter) were purchased from The Jackson Laboratory. *Manf* ^fl/fl^ mice and *Pcp2-Cre*^+/-^ mice were crossed for two generations to generate *Manf* ^fl/fl^; *Pcp2-Cre*^+/-^ mice which have specific MANF knockout in the PCs (referred to as KO and thereafter). Their littermates that were *Manf* ^fl/fl^; *Pcp2-Cre*^-/-^ were used as control animals (referred to as control and thereafter). All mice were genotyped by polymerase chain reaction (PCR) analysis using the Fast Tissue/Tail PCR Genotyping Kit (G1001, EZ BioResearch) with primers (Integrated DNA Technologies) as follows: Manf (forward) 5’-TGAAGCAAGAGGCAAAGAGAATCGG-3’, Manf (reverse) 5’-TGCTCAGCTGCAGAGTTAGAGTTCC-3’; Cre (forward) 5’-GGTTCGCAAGAACCTGATGG-3’, Cre (reverse) 5’-GCCTTCTCTACACCTGCGG-3’. PCR products were visualized by electrophoresis on 1% agarose gel and stained with ethidium bromide (Thermo Fisher Scientific).

### Alcohol Administration

A binge alcohol exposure paradigm was used for this study as it has been widely used to produce alcohol neurotoxicity in rodents, mimicking the neuropathological conditions in human with alcohol use disorders [57]. Both male and female adult mice at the age of four to five months old received equal volume of H2O or ethanol (200 proof, Decon Labs, Inc, Swedeland, PA) (5 g/kg, 25% ethanol w/v) via intragastric gavage once daily for 10 days. The resulting blood alcohol concentration (BAC) peaks above 300 mg/dl one hour after the last ethanol administration [28]. All mice were given ad libitum access to food and water throughout the period of alcohol administration.

### Behavioral Tests

Behavioral tests were performed 10 days after the last alcohol exposure. A total of 147 animals (74 males, 73 females) were used for behavior tests, with at least 8 animals in each treatment group for all the tests. All mice were given ad libitum access to food and water throughout the tests.

#### Open Field

The open field (OF) test is a common behavioral assay that measures locomotor activity in rodents [58]. Before the test, animals were removed from their home cage and brought to the testing room in a clean holding cage to acclimate for 10 min as a habituation period. The test was conducted 15 minutes for each animal in a multi-unit open field arena measured 50 cm × 50 cm × 40 cm (San Diego Instruments, San Diego, CA). The total distance travelled was recorded using the EthoVision XT 17 video-tracking software (Noldus Information Technology, Leesburg, VA, USA).

#### Rotarod Test

The rotarod test is a commonly used method to evaluate the motor coordination deficits in rodents using an animal’s natural avoidance of fall. Mice were placed on a rotating rod 18” above the ground in the rotarod apparatus (San Diego Instruments, San Diego, CA). The latency to fall, which is the accumulative time for each mouse to remain on the rod, was recorded automatically with infrared sensors as an indicator for motor coordination. Mice were first trained on the rotating rod at a constant speed of 4 rpm for 60 seconds. Then the rod was rotated from a speed of 4 rpm to 40 rpm over the course of 300 seconds. Three consecutive 300 seconds trials were performed for each mouse, and the latency to fall was recorded, in which 300 seconds were set as the maximum time. The mean of the 3 trials was calculated for each mouse.

#### Balance Beam Test

Balance beam test was used to assess motor coordination. Mice were placed at the end of an 80 cm-long wood beam elevated 50 cm from the floor. An open cage was attached to the other end of the beam. During the training phase, each animal was allowed to cross a beam (3.0 cm wide) freely twice. After training, mice were required to cross the beams with various width (2.0 cm, 1.0 cm, and 0.5 cm) from one end to the other end in the open cage. The time spent for the crossing was recorded for each animal. If an animal stopped crossing the beam, its tail was gently touched to encourage movement. If an animal fell from the beam, it was placed back on the end of the beam to restart crossing.

#### Three-chamber Sociability Task

At least 30 minutes before testing starts, all animals were acclimated in the testing room. A matte black plastic box arena with three chambers and openings between the chambers was custom made for the test. The two side chambers each contains a plastic cylinder with holes allowing interaction. Each animal was habituated in the arena with empty cylinders for 10 minutes. Then it is followed with 10 minutes testing session, during which a novel mouse was placed in one chamber’s cylinder and a novel object was placed in the other chamber’s cylinder. Novel mice were wild type C57BL/6 mice of the same sex and age that the tested mice had never been exposed to previously. The total distance travelled, and the time spent in each chamber and with each cylinder were recorded and analyzed by the EthoVision XT 17 system.

#### Barnes Maze

The Barnes maze is a white circular platform with a diameter of 36’’ and elevated 40’’ from the floor. It has 20 holes with a diameter of 2’’ that are equally distributed around the perimeter edge (San Diego Instruments w/ Custom Legs, San Diego, CA). The test lasted for 6 days. On day 0, each mouse was placed on the center of the maze to habituate on the maze and explore the maze freely. An escape box was placed under one of the holes that was not to be used during training and probe testing trials. All other holes were blocked. After 1 minute, mouse was gently directed into the escape box. Day 1-4 were for training trials to test the mice’s ability in learning. There were 4 visual cues present on the walls and the escape hole location was rotated between mice. Each mouse was placed on the center of the maze and explore the maze to find and enter the escape box. The trial ended when the mouse entered the escape box or 3 minutes have elapsed. Training was conducted 4 times per day for 4 consecutive days. Mice were rested 10-15 minutes between trials. On day 5, mice were given a 1.5-minute probe testing trial to test for short-retention memory. All holes on the maze were blocked, including the target hole. Each mouse was placed on the center of the maze to explore and visit the location where the escape box used to be. Three parameters were recorded during training and testing trials, including the primary distance which is the total distance traveled by mice to find the escape hole, the primary latency which is the time that an animal took to encounter the escape hole for the first time, and the search strategies which are defined as random (animals move randomly across the platform until the escape box is found), serial (animals travel through consecutive holes around the periphery of the maze until they find the escape box), and direct (animals navigate directly to the correct quadrant and escape box without crossing the maze center more than once and with less than two errors). The time spent in the target quadrant during probe testing trials was also recorded. Data was analyzed using the EthoVision XT 17 system.

### Immunohistochemistry and Immunofluorescence

Mice were anesthetized with intraperitoneal injection of ketamine/ xylazine solution (100 mg/kg/10 mg/kg, Butler Schein Animal Health, Portland, ME), and then perfused intracardially with PBS followed by 4% paraformaldehyde (PFA, 15714, Electron Microscopy Sciences) in PBS (pH 7.4). Cerebella were collected, post-fixed in 4% PFA in PBS for 48 h, and then paraffin embedded in the Comparative Pathology Laboratory at University of Iowa Department of Pathology. Cerebellar were sectioned through the midline vermis at the thickness of 5 µm using a rotary microtome (Leica), mounted onto superfrost/plus slides (48311-703, VWR, West Chester, PA), and processed for immunohistochemistry (IHC) or immunofluorescence (IF) staining as previously described [30]. After deparaffinization, slides were antigen retrieved in 10 mM sodium citrate buffer (pH 6.0) in a high pressure cook for 7 minutes, blocked with 1% BSA/2% goat serum/0.1% Triton x-100 at room temperature for 1 hour, and incubated with primary antibodies at 4 °C overnight. For IHC, slides were treated with 0.3% H2O2 in methanol for 15 minutes, then incubated with HRP-conjugated biotinylated secondary antibodies (1:200, Vector Laboratories Inc, Newark, CA) for 1 hour and incubated with Avidin-biotin-peroxidase complex (ABC) (Vector Laboratories Inc). The immunolabeling was visualized using DAB peroxidase (HRP) substrate kit (SK-4100, Vector Laboratories Inc) and imaged using Olympus BX81 microscope (Olympus, Waltham, MA). For IF, slides were incubated with Alexa fluor-conjugated secondary antibodies (1:200, Life Technologies, Carlsbad, CA) in the dark for 1 hour, and counterstained with DAPI (D9542, Sigma-Aldrich, St. Louis, MO). Slides were sealed with VECTASHIELD mounting medium (H-1400, Vector Laboratories Inc) and imaged using the Olympus IX81 microscope (Olympus).

The following antibodies were used for this study: anti-ARMET/ARP (MANF) (ab67271) and anti-ATF6 (ab203119) were from Abcam (Cambridge, MA); anti-Calbindin-D-28K (C9848) was from Sigma-Aldrich (St. Louis, MO); anti-GRP78 was from Novus Biologicals (Littleton, CO); anti-β-Actin (CST3700), anti-phosphorylated eIF2α (CST3398), anti-Calbindin (CST13176), and anti-cleaved Caspase-3 (CST9661) were from Cell Signaling Technology (Danvers, MA); anti-XBP1s (658802) was from BioLegend (San Diego, CA). HRP-conjugated anti-rabbit (GENA934) and anti-mouse (GENA931) secondary antibodies were from GE Healthcare Life Sciences (Piscataway, NJ). Alexa-488 conjugated anti-mouse (A21202), Alexa-594 conjugated anti-mouse (A11005) and Alexa-594 conjugated anti-rabbit antibodies (A11012) were from Life Technologies (Carlsbad, CA).

### Stereological Cell Counting

Every twentieth section from the serial sagittal sections across the cerebellar vermis was immunolabeled with anti-Calbindin-D-28K for PCs. The number of PCs in cerebellar lobule II of each section were counted by unbiased stereology that uses systematic random sampling method for counting. The Optical Fractionator probe was performed on an Olympus BX-51 microscope equipped with ASI motorized stage with XYZ encoder. Stereo Investigator software (v11, MBF Bioscience, Williston, VT) was used to estimate the total number of PCs in lobule II of each brain sample. The contours were drawn manually to mark the PC layer. Based on our previous experience, we used a sampling grid of 800×800 μm and a counting frame of 200×200 μm for all the brain samples measured, which generated the coefficient of error (CE) of 0.07 approximately. We counted the number of PCs at a final magnification of 100x, using the unbiased counting rule that uses two exclusion edges (left and lower) and two inclusion edges (upper and right) on the images while counting.

### Immunoblotting

Protein was extracted from mice brains as described previously [48]. Protein concentration was determined using the DC protein assay according to the manufacture’s instruction. For immunoblotting, 30 µg protein samples were separated on 12% polyacrylamide gels by electrophoresis and transferred to nitrocellulose membranes and blocked in 5% BSA/1xTBS/0.05% Tween-20 for 1 h at room temperature prior to incubation with primary antibodies at 4°C overnight. Subsequently, membranes were washed with TBST and incubated with secondary antibodies conjugated to horseradish peroxidase. Blots were developed using the Cytiva Amersham ™ ECL™ Prime Western Blotting Detection Reagent on Chemi™Doc imaging system (Bio-Rad Laboratories, Hercules, CA). Band intensity was quantified using Image Lab software (Bio-Rad Laboratories).

### TUNEL Assay

TUNEL assay was performed using the In Situ Cell Death Detection Kit Fluorescein (11684817910, Roche, Indianapolis, IN) according to the manufacturer’s instructions. Slides were deparaffinized and treated with 10 mM sodium citrate buffer (pH 6.0) in a high pressure cook for 7 minutes. Then, slides were incubated with TUNEL reaction mixture at 37 °C in a humid chamber in the dark for 60 minutes. Slide incubated in labeling solution without terminal transferase was used as the negative control and slide digested with DNase I and incubated with TUNEL reaction mixture was used as the positive control. The slides were then counterstained with DAPI and imaged with the Olympus IX81 inverted fluorescent microscope (Olympus).

### Visium Spatial Transcriptomics Analysis

STA is a next-generation whole transcriptome analysis measuring total mRNA in tissue sections while maintaining the spatial attributes of cells. It is a powerful tool to dissect gene expression heterogeneity in the architecture of intact tissue sections [59]. Tissue processing for Visium spatial transcriptomics (Visium Gene Expression Slide & Reagent Kit, PN-1000184, 10x Genomics, Pleasanton, CA) was performed in the Iowa Neuroscience Institute (INI) NeuroBank Core following the instruction of the tissue preparation guide demonstrated protocol CG000240. Twelve fresh mouse brains (male control n=3, male KO n=3, female control n=3, female KO n=3) were dissected and snap frozen in a metal beaker with cooled isopentane in a foam dewar with liquid nitrogen bath. A small piece of the cerebral cortex of each frozen brain was used for RNA extraction. RNA quality was assessed in the Iowa Institute of Human Genetics (IIHG) Genomics division using RNA Integrity Number (RIN) and all samples were confirmed with a RIN above 7. Cerebellum from the 12 samples were cryosectioned through the midline vermis into 10μm sections. One section of each cerebellum was mounted onto the Visium spatial slides (2000233, 10x Genomics) within the frames of capture areas. Slides were then processed with methanol fixation, H&E staining and imaging following demonstrated protocol CG000160. Prior to library preparation, the optimal permeabilization condition was determined following the tissue optimization demonstrated protocol CG000238. A total of 18 minutes of permeabilization was chose based on the optimalization result. After reverse transcription and second strand synthesis, the amplified cDNA samples from the Visium slides were transferred, purified, and quantified for library preparation. Sequencing libraries were prepared by the IIHG Genomics Division, according to the Visium Spatial Gene Expression Reagent Kits User Guide CG000239. The molar concentrations of the indexed libraries were measured using the 2100 Agilent Bioanalyzer (Agilent Technologies, Santa Clara, CA) and combined in a ratio to achieve at least 50,000 sequencing read pairs per tissue covered spot. The concentration of the pool was determined using the Illumina Library Quantification Kit (KAPA Biosystems, Wilmington, MA) and sequenced on the Illumina NovaSeq 6000 genome sequencer using the 28 bp Read 1, 10 bp i7 index, 10 bp i5 index, and 90 bp Read 2 run configuration using SBS v1.5 sequencing chemistry.

Bioinformatic analysis of the Visium data was carried out by the IIHG Bioinformatics division. Raw FASTQ files and H&E images were processed with Space Ranger (10X Genomics, version 1.3.0) using the mm10-2020-A reference transcriptome. Quality control (QC) showed no quality concerns for all samples. The resulting data matrices were subsequently imported into R (version 4.0.0) and analyzed using the Seurat package (version 4.0.5). Gene expression was normalized and scaled using the SCTransform function. Based on visual inspection of the elbow plot, the first 30 Principal Components (PC) were used in UMAP-based dimensional reduction. The FindClusters function, with a resolution of 0.5, was then used to assign cells to clusters. After confirming that the cell clusters were anatomically accurate by visual inspection, we used the Seurat cluster assignments to create pseudobulk RNA-Seq samples. This aggregation of counts for each individual cluster was done using the aggregateBioVar package (version 0.99.2). These counts were then normalized and transformed using a variance stabilizing transformation (vst) and principal components analysis (PCA) was performed to visualize sample clusters. For statistical analysis of the data, the DESeq2 package (version 1.30.1) was used. In brief, a model incorporating all the experimental factors was created and Wald tests were used to compute statistical metrics. A gene was considered to have a statistically significant change in expression if the False Discovery Rate (FDR) was less than 10% (adjusted *p* value <0.1). A gene was considered to have biologically significant change in expression if the base mean was larger than 5 and the |log2Fold Change (FC)| was larger than 0.2. Genes found to be differentially expressed under these criteria were used as input for enrichment analysis (iPathwayGuide, Advaita Bioinformatics, Ann Arbor, MI). Visium spots overlay with image were visualized using Loupe browser (10x Genomics, version 8.1.2). Results from the statistical analysis were visualized using heatmaps (created with pheatmap, version 1.0.12). Venn diagram was generated using the free online tool https://bioinformatics.psb.ugent.be/webtools/Venn/.

### Xenium In Situ Analysis

Xenium is a probe-based high throughput in situ spatial RNA profiling platform. It allows effective detection for up to 5,000 genes on the same tissue section simultaneously [60]. Cerebella from twelve animals (male control n=3, male KO n=3, female control n=3, female KO n=3) were collected, fixed, and embedded in paraffin as described in the immunohistochemistry and immunofluorescent section. Formalin-fixed paraffin-embedded (FFPE) tissue blocks were sent to the Advanced Genomics Core at the University of Michigan for processing using 10x Genomics demonstrated protocols CG000578, CG000580, CG000582, and CG000613. Tissue sections (5µm) were collected onto Xenium slides (3000941, 10x Genomics) where sections were deparaffinized and decrosslinked. RNA within the tissue was labeled using circularizable DNA probes targeting 300 genes in a standalone custom gene panel (1000648, 10x Genomics, design ID: HNGXTA). Following ligation, the DNA probes were enzymatically amplified followed by autofluorescence quenching and nuclei staining with DAPI. Xenium slides were loaded into the Xenium Analyzer instrument for imaging and analysis, where fluorescently labeled oligos bound the amplified DNA probes, and samples underwent successive rounds of fluorescent probe hybridization, imaging, and removal to generate optical signatures which were converted into a gene identity. Following the Xenium run, slides were H&E stained and high-resolution images captured for inclusion in the final data set. Acquired data was transferred off the Xenium instrument and data visualized using the Xenium Explorer (10x Genomics, version 3.2.0) and third-party analysis tools.

Bioinformatic analysis of the Xenium data was carried out by the Genomics Research and Technology Hub at University of California. Xenium data was downloaded and imported using R package Seurat (version 5.0.3) LoadXenium function. Cell barcodes for regions of interest were obtained from Xenium Explorer and used to subset the raw gene count matrix. Seurat objects for each of the 12 samples were then created and merged ensuring that each gene was expressed in at least 3 cells. After normalization, highly variable genes were then identified using FindVariableFeatures function, and dimension reduction with principal component analysis (PCA) was performed using RunPCA function and uniform manifold approximation and projection using RunUMAP function. To identify differentially expressed genes between different conditions, Wilcoxon rank sum test was used with FindMarkers function. Pseudobulk analysis was performed using DESeq2 (version 1.42.0) where raw counts of cells from the same sample were summed and normalized within DESeq2. Differential gene expression analysis was performed using DESeq2 built in generalized linear models to compare the KO vs control samples. A gene was considered to have a statistically significant change in expression if the False Discovery Rate (FDR) was less than 10% (adjusted *p* value <0.1). A gene was considered to have biologically significant change in expression if the base mean was larger than 5 and the |log2FC| was larger than 0.2. Volcano and violin plots were generated using GraphPad Prism 10. The enrichment analysis was performed using g:Profiler (version e112_eg59_p19_25aa4782) with g:SCS multiple testing correction method applying significance threshold of 0.05 [61].

### Statistical Analysis

All statistical analyses were performed using the GraphPad Prism 10 software unless notified specifically. Data were expressed as mean ± SEM. Differences among experimental groups were analyzed by Student t-test, one-way ANOVA or two-way ANOVA with *p*<0.05 being considered statistically significant. In cases where significant differences were detected, specific comparisons between treatment groups were examined with the Tukey’s post-hoc test.

## Results

### Characterization of PC specific MANF KO mouse model

MANF is strongly expressed in the developing and adult mouse PCs. Low levels of MANF expression can be detected in the developing PCs at postnatal day (PD) 4. It was increased by PD 7 and expressed strongly in the 4 months old adult cerebellum PCs (Figure S1A). Immunoblotting of MANF using the 1- and 4-month mouse cerebellum lysates demonstrated the expression of MANF in both male and female cerebellum, with female expressing higher levels of MANF than male (Figure S1B and S1C). The expression of Cre recombinase in the *Pcp2-Cre* line is driven by the promoter of the mouse *Purkinje cell protein-2* (*Pcp2*) gene that is expressed exclusively in PCs in the brain, thus it is widely used to target gene deletion in PCs [62, 63]. The *Pcp2-Cre* line exhibits onset of Cre recombinase in PCs around PD 6-7 and is fully established by PD 14-15 and remains constant throughout the lifespan of the mice [63, 64]. To confirm MANF knockout in PCs, brains from age matched control (*Manf* ^fl/fl^) and KO (*Manf* ^fl/fl^; *Pcp2-Cre*^+/-^) mice at PD 7, 14, and 21 were immunolabeled with anti-MANF antibody and PC marker anti-Calbindin antibody. At PD 7, MANF expression was observed in both control and KO PCs, and reduced expression of MANF in KO PCs was not detected yet (Figure 1A). At PD 14, MANF was significantly reduced in the KO PCs (Figure 1B). By PD 21, MANF expression was completely absent in the KO PCs (Figure 1C). We examined the body weight and brain weight of control and KO mice. Regardless of gender and genotype, body weight of control and KO mice were comparable at PD 7 and PD 21 (Figure 1E). Control and KO male and female mice were raised to adulthood (Figure 1D). By 4 months and 1 year’s old, male mice were significantly heavier than female mice (Figure 1E), but no difference in body weight and brain weight was observed between control and KO (Figure 1E and 1F). In the adult cerebellum, MANF remained to be absent in the KO PCs in both male and female (Figure 1G). MANF deficiency was specific to PCs as its expression in the sporadic interneurons in the granular layer was not affected (Figure 1H, arrows) and its expression in other brain regions such as in the cerebral cortex, hippocampus, and olfactory bulb was not affected neither (Figure S2).

**Figure 1.**
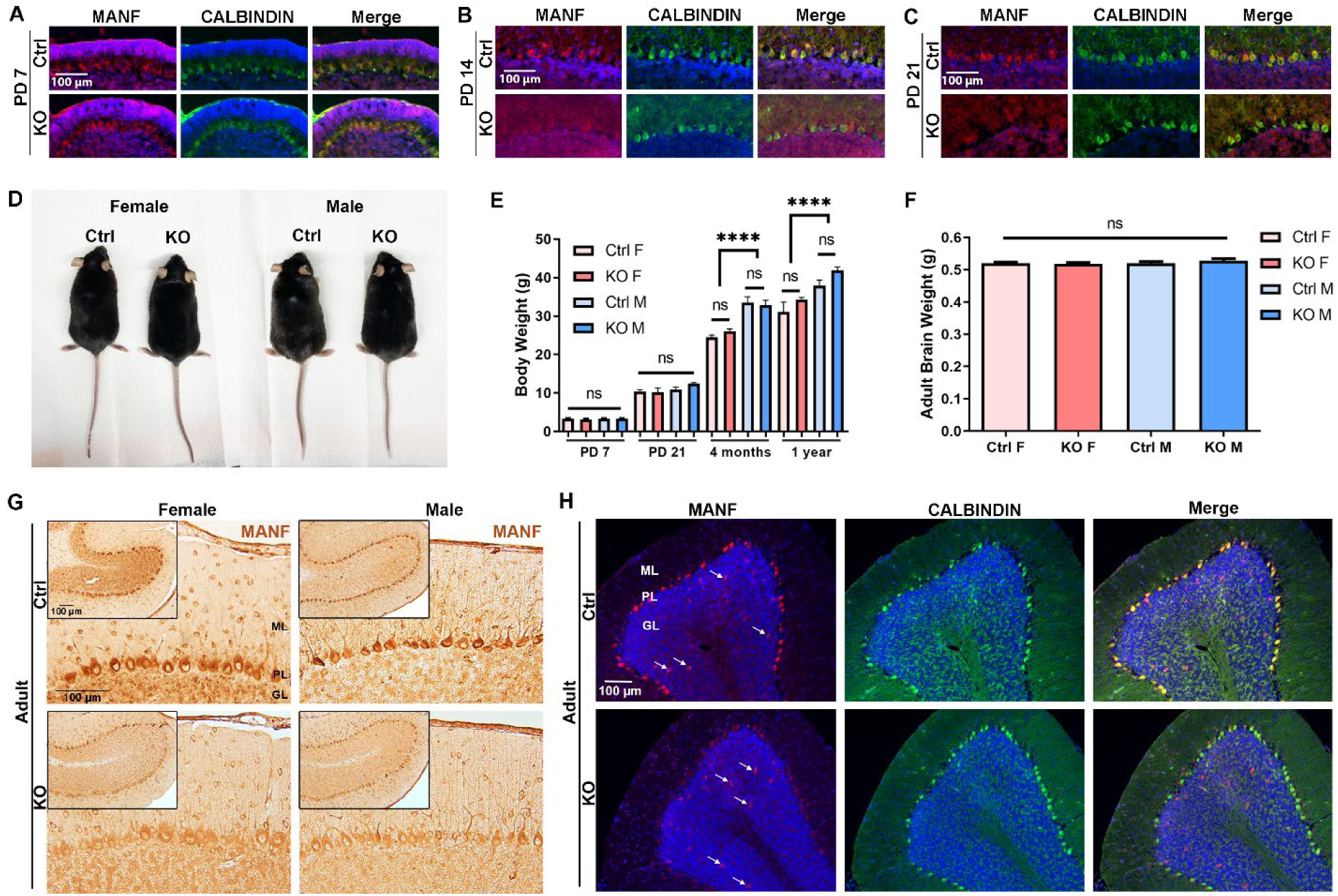
Characterization of PC specific MANF KO mouse model. **A-C.** Representative immunofluorescent images of MANF (red) and PC marker Calbindin (green) in the control and KO cerebellum at postnatal day (PD) 7 (A), PD 14 (B), and PD 21 (C). **D-F.** Morphology (D), body weight (E), and brain weight (F) of the adult control and MANF KO mice. The data was expressed as mean ± SEM. n=8-16 per group for body weight; n=6 per group for brain weight. Two-way ANOVA followed by Tukey’s *post hoc* test. ****p < 0.0001; ns not significant. **G.** Representative immunohistochemistry images of MANF in the adult control and KO cerebellum. Insets show the entire cerebellum lobule II. **H.** Representative immunofluorescent images of MANF (red) and PC marker Calbindin (green) in the adult control and KO cerebellum. Arrows indicated MANF expression in cells located in the granular layer. ML, molecular layer; PL, Purkinje cell layer; GL, granule cell layer.

### MANF deficiency exacerbates binge alcohol exposure induced motor deficits

Control and KO male and female mice at the age of 4-5 months were exposed with either water or ethanol (5g/kg/day) daily for 10 days through intragastric gavage and their motor function was evaluated in open field, balance beam, and rotarod tests 10 days after the last gavage (Figure 2). The body weight of all animals was recorded during gavage and behavior tests. Although both male and female control and KO mice showed a significant body weight loss due to ethanol gavage, it recovered to pre-gavage level by the time of behavior tests (Figure S3). For open filed test, the total distance traveled was recorded. In females, main effects of genotype and ethanol were found with KO females travelled less distance than control females and ethanol treated animals travelled less than control groups (Figure 3A). In addition, an effect of genotype and ethanol interaction was observed. Post hoc analysis indicated that ethanol exposed KO female traveled significantly less distance than all the other groups (Figure 3A). In males, only a main effect of genotype was observed with KO males travelled less distance than control males, while ethanol showed no effects (Figure 3B). Post hoc analysis showed that ethanol exposed KO males traveled less than water treated control males (Figure 3B). Animals were then tested in rotarod test. In females, a main effect of genotype and an interaction of genotype and ethanol treatment were observed. Post hoc analysis indicated a significantly reduced latency to fall from the rotarod for ethanol treated KO females when compared to all the other groups (Figure 3C). No difference was observed for males in rotarod test (Figure 3D). Lastly, animals were tested in balance beam test. In females, although no differences were detected when the beam width was 2.0 and 1.0 cm, there were significant main effects of genotype and ethanol, and an effect of genotype and ethanol interaction when the beam width was as narrow as 0.5 cm (Figure 3E). Ethanol exposed KO female spent significantly longer time to cross the 0.5 cm beam when compared to other treatment groups (Figure 3E). For males, there was a main effect of genotype that KO males spent longer time to cross the beam than control males regardless of the beam widths. No main effect of ethanol was observed in male balance beam test, although when the beam width is at 0.5 cm, post hoc analysis found a significant increased crossing time when compared ethanol treated KO males with control or ethanol treated control males (Figure 3F). These results from open field, rotarod, and balance beam tests indicated that PCs MANF deficiency impaired motor functions. Binge alcohol exposure exacerbated the motor deficits in mice with PC-specific MANF deficiency, and female KOs were more sensitive than male KOs to alcohol induced motor impairments.

**Figure 2.**
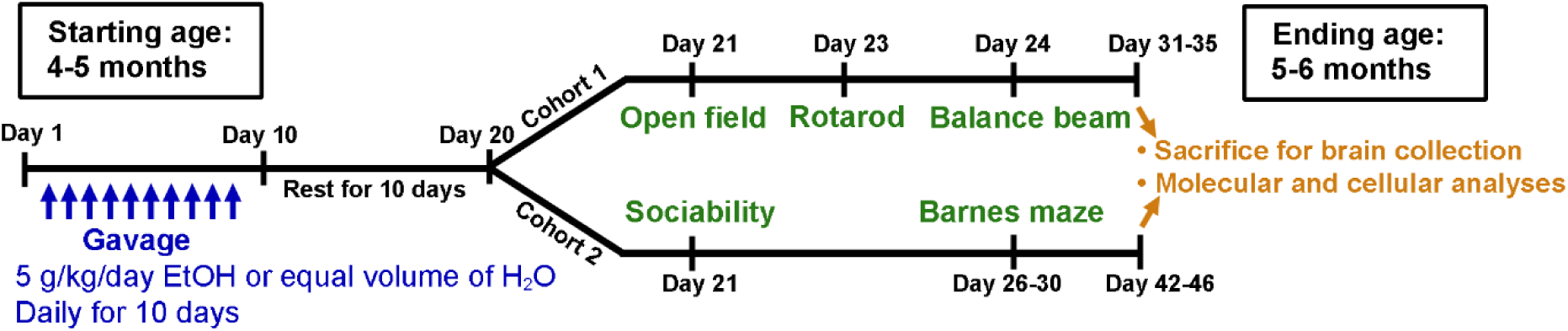
Schematic illustration of the experimental timeline. Both male and female adult mice at the age of four to five months old received equal volume of H2O or ethanol (5 g/kg, 25% ethanol w/v) via intragastric gavage once daily for 10 days. Behavioral tests were performed 10 days after the last ethanol exposure. Cohort 1 animals were tested in open field, rotarod, and balance beam tests. Cohort 2 animals were tested in 3-chamber sociability task and Barnes maze test. After the tests, animals were euthanized and brains collected for further cellular and molecular analysis.

**Figure 3.**
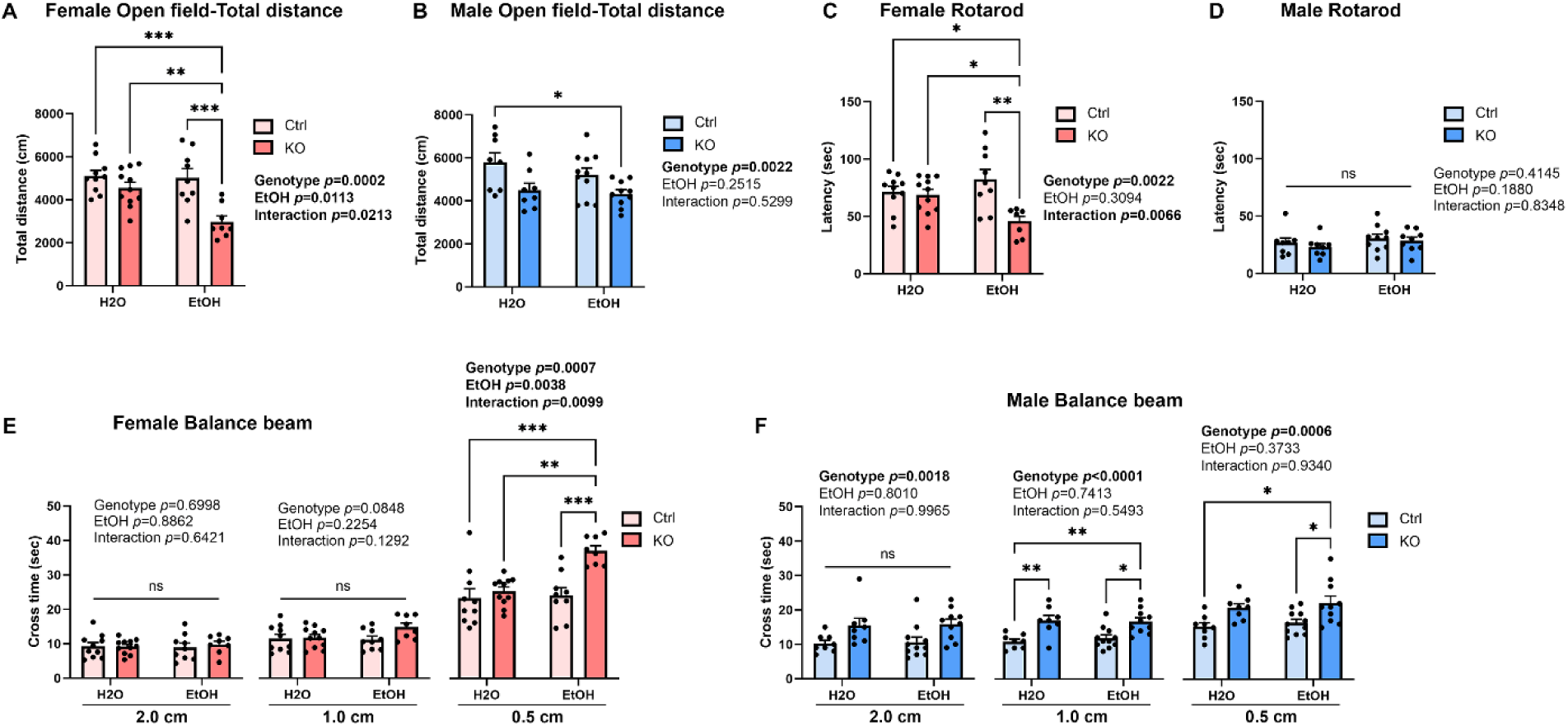
Effects of ethanol exposure on motor functions in PC specific MANF KO mice. **A-B.** Total distance traveled in open field test for water- and ethanol-exposed control and KO female (A) and male (B) mice. **C-D.** The latency to fall from the accelerating rotarod in water- and ethanol-exposed control and KO female (C) and male (D) mice. **E-F.** The time to cross the balance beam in water- and ethanol-exposed control and KO female (E) and male (F) mice. All data was expressed as mean ± SEM. n=8-11 per group. Two-way ANOVA followed by Tukey’s *post hoc* test. Significant main effect *p* values were highlighted in bold; \**p*< 0.05; \*\**p*<0.01; \*\*\**p*<0.001; ns not significant.

### MANF deficiency does not affect binge alcohol exposure induced alteration in social interaction

Binge alcohol drinking is well known to influence sociability in both human and mice [65, 66]. Dysfunction of cerebellum and PCs have also been implicated in social behavior impairments [67–69]. To test whether social behavior was altered by alcohol and PCs MANF deficiency, animals were examined in the 3-chamber sociability test. Female mice stayed significantly longer time in the social chamber than the object chamber (Figure 4A) and spent significantly longer time interacting with the social cylinder than the object cylinder (Figure 4C), indicating a preference in social interaction. More importantly, a main effect of ethanol was observed with ethanol treated females showing reduced social cylinder time than water treated females. Post hoc analysis found a significant reduction of social cylinder time in ethanol treated KO females when compared with water treated controls (Figure 4C). For males, although no chamber preference was observed (Figure 4B), males still exhibit a significant preference of social cylinder over object cylinder (Figure 4D), indicating that males also prefer social interactions. However, no effect of ethanol nor PCs MANF KO on sociability was observed in male. These results indicated that alcohol impaired social interaction behaviors in the female but not male mice, and it was not affected by PCs MANF deficiency.

**Figure 4.**
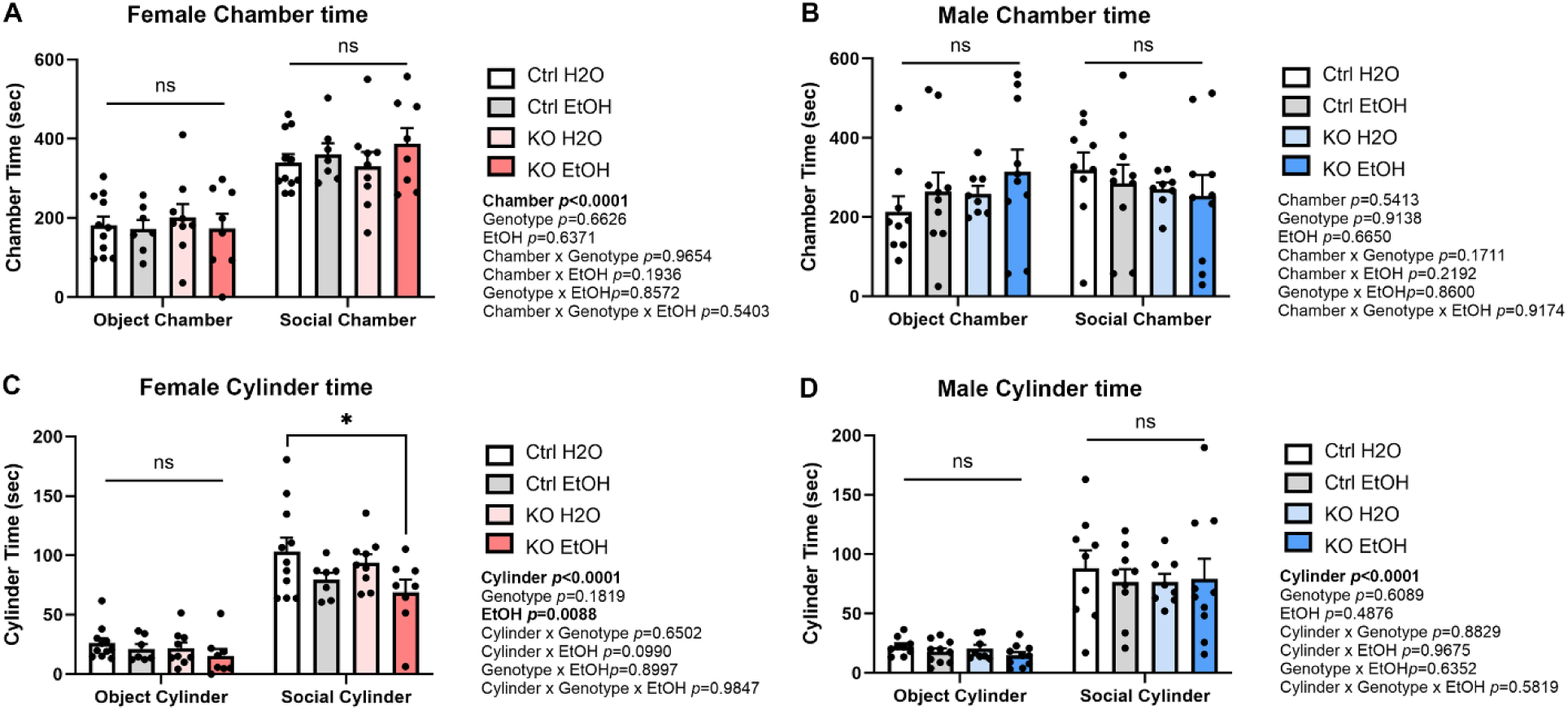
Effects of ethanol exposure on social behaviors in PC specific MANF KO mice. **A-B.** Time spent in the object and social chambers in water- and ethanol-exposed control and KO female (A) and male (B) mice. **C-D.** Time interacting with the object and social cylinders in water- and ethanol-exposed control and KO female (C) and male (D) mice. All data was expressed as mean ± SEM. n = 8–11 per group. Data was analyzed by three-way ANOVA followed by Tukey’s *post hoc* test. Significant main effect *p* values were highlighted in bold; \**p*<0.05; \*\**p*<0.01; ns not significant.

Animals were also accessed for their cognitive function in learning and memory, but no effect of alcohol nor MANF KO was observed (Figure S4 and S5).

### Alcohol induces PC degeneration in MANF deficient females

After the behavior tests, brains from all animals were collected to investigate the cellular and molecular effect of MANF deficiency and alcohol neurotoxicity on PCs. Since the anterior portion of the cerebellum vermis (lobule 1-IV) is most sensitive to alcohol neurotoxicity [18, 70, 71], we choose to focus on the cerebellar lobule II for cellular and molecular analysis to avoid variations among different lobules. Tissue sections through the cerebellar vermis were immunolabeled with PC specific marker Calbindin to reveal the number and morphology of PCs. Stereological cell counting found a significant reduction in the number of PCs in ethanol treated KO females (Figure 5A and 5C). The remaining PCs in ethanol treated female KO cerebellum showed a significant decrease in cell body size, indicating PC shrinkage and degeneration (Figure 5E). Interestingly, we also observed a change in the distribution of Calbindin in the ethanol treated MANF KO PCs. Calbindin normally exhibits a diffused expression across the cytoplasm, dendrites, axon, and nucleus in PCs, but in ethanol treated female KO PCs, there was a diminished perikaryal staining accompanied with predominant intranuclear immunoreactivity (Figure 5A, insets). In males, however, neither ethanol treatment nor MANF deficiency affected the PC number, cell size, or Calbindin distribution (Figure 5B, 5D, and 5F).

**Figure 5.**
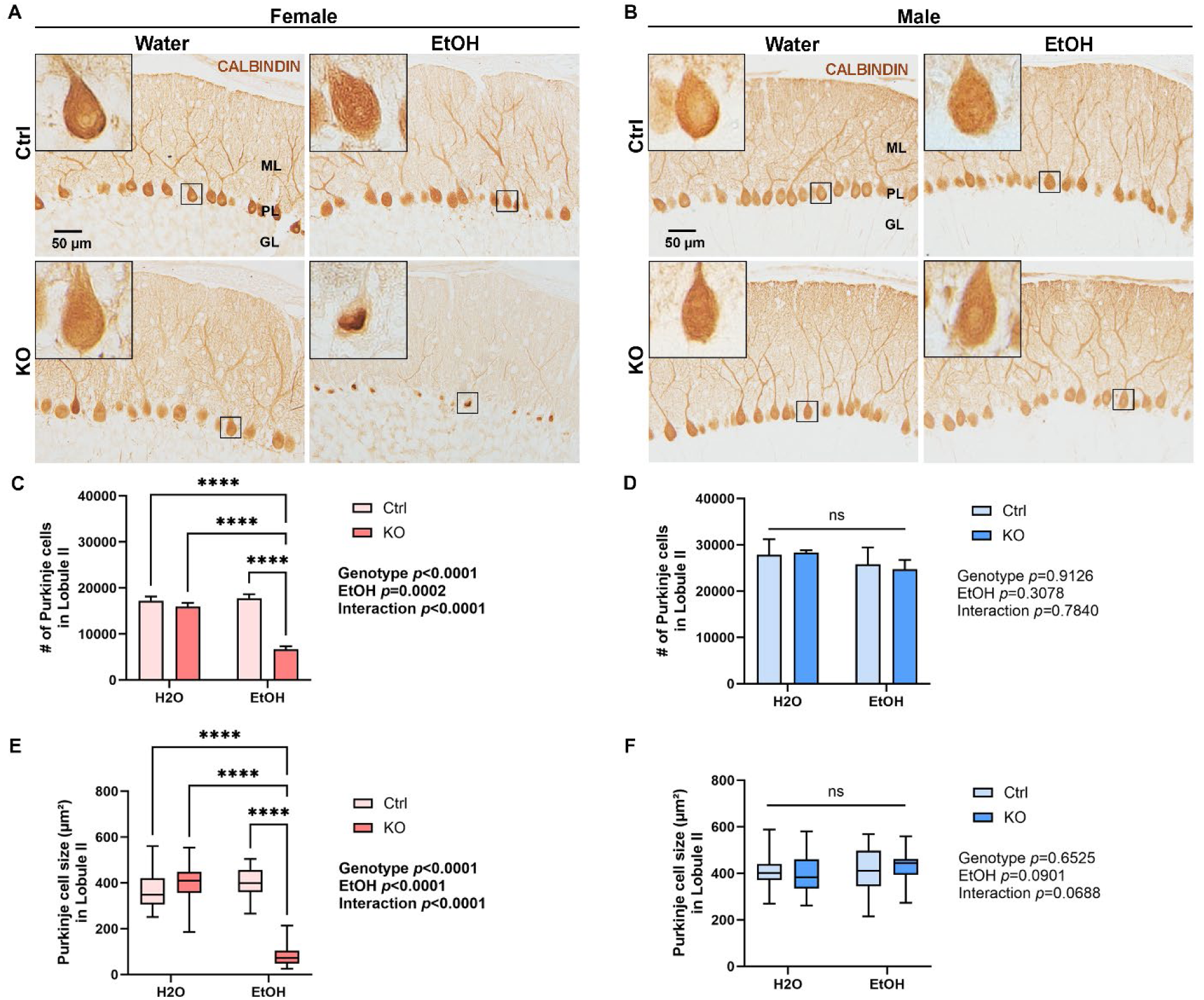
Female MANF deficient PCs are more sensitive than males to ethanol induced morphological changes. **A-B.** Representative immunohistochemistry images of PC marker Calbindin in the adult control and KO cerebellum in female (A) and male (B) after water- or ethanol-exposure. Insets represent the enlarged view of the PCs. Note the diminished perikaryal staining and predominant intranuclear immunoreactivity of Calbindin in the ethanol treated female KO PCs. **C-F.** Quantification of the number (C, D) and size (E, F) of PCs in female (C, E) and male (D, F) cerebellum lobule II. The data was expressed as mean ± SEM. n=4 per group. Two-way ANOVA followed by Tukey’s *post hoc* test. Significant main effect *p* values were highlighted in bold; \*\*\*\**p*<0.0001; ns not significant.

### Alcohol induces ER stress and apoptotic cell death in female MANF deficient PCs

PCs are particularly vulnerable to alcohol induced disruption in ER homeostasis [7, 8]. We have previously reported that neuronal MANF deficiency exacerbated alcohol induced ER stress in the brain [31, 32]. To examine if alcohol can cause ER stress in MANF deficient PCs, tissue sections through the cerebellar vermis were immunolabeled with MANF interacting ER molecular chaperon proteins GRP78 and HYOU1, and UPR proteins XBP1s, p-PERK, p-eIF2α, and ATF6. Results in female showed a significant upregulation of all the markers in ethanol treated MANF deficient PCs (Figure 6A-6F). On the contrary, the expression of most markers was not affected by ethanol or MANF deficiency in male PCs, except for GRP78 that it was significantly upregulated in ethanol treated male MANF deficient PCs (Figure 7A-7F).

**Figure 6.**
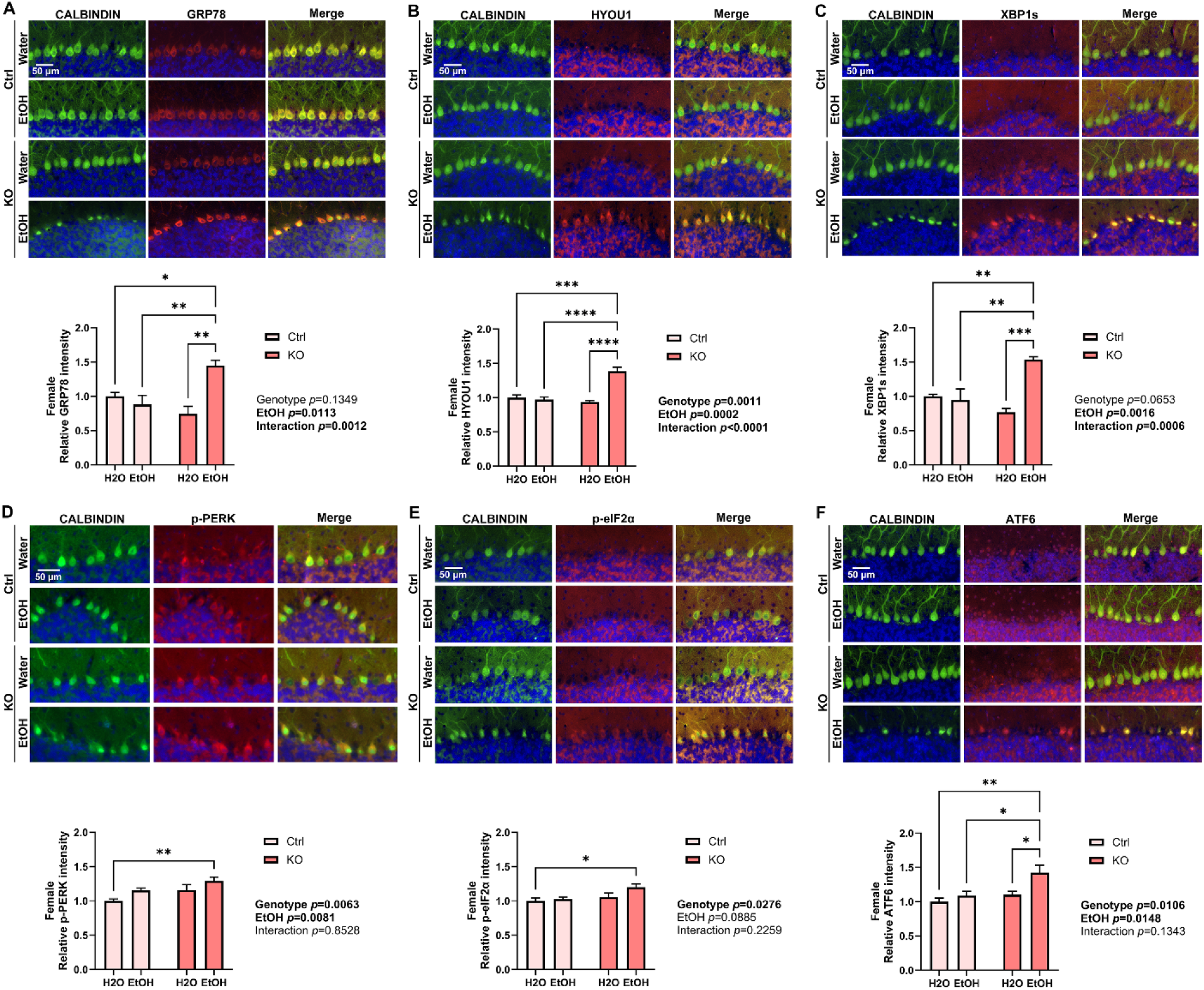
Ethanol exposure induces ER stress in female MANF KO PCs. **A-F.** Representative immunofluorescent images and quantifications for the expression of ER stress markers GRP78 (A), HYOU1 (B), XBP1s (C), p-PERK (D), p-eIF2α (E), and ATF6 (F) in water- and ethanol-exposed control and KO female mice cerebellum. PC marker Calbindin was co-labeled in green. Quantification of ER stress markers fluorescence intensity was expressed as mean ± SEM. n=3-4 per group. Two-way ANOVA followed by Tukey’s *post hoc* test. Significant main effect *p* values were highlighted in bold; \**p*< 0.05; \*\**p*<0.01; \*\*\**p*<0.001; \*\*\*\**p*<0.0001; ns not significant.

**Figure 7.**
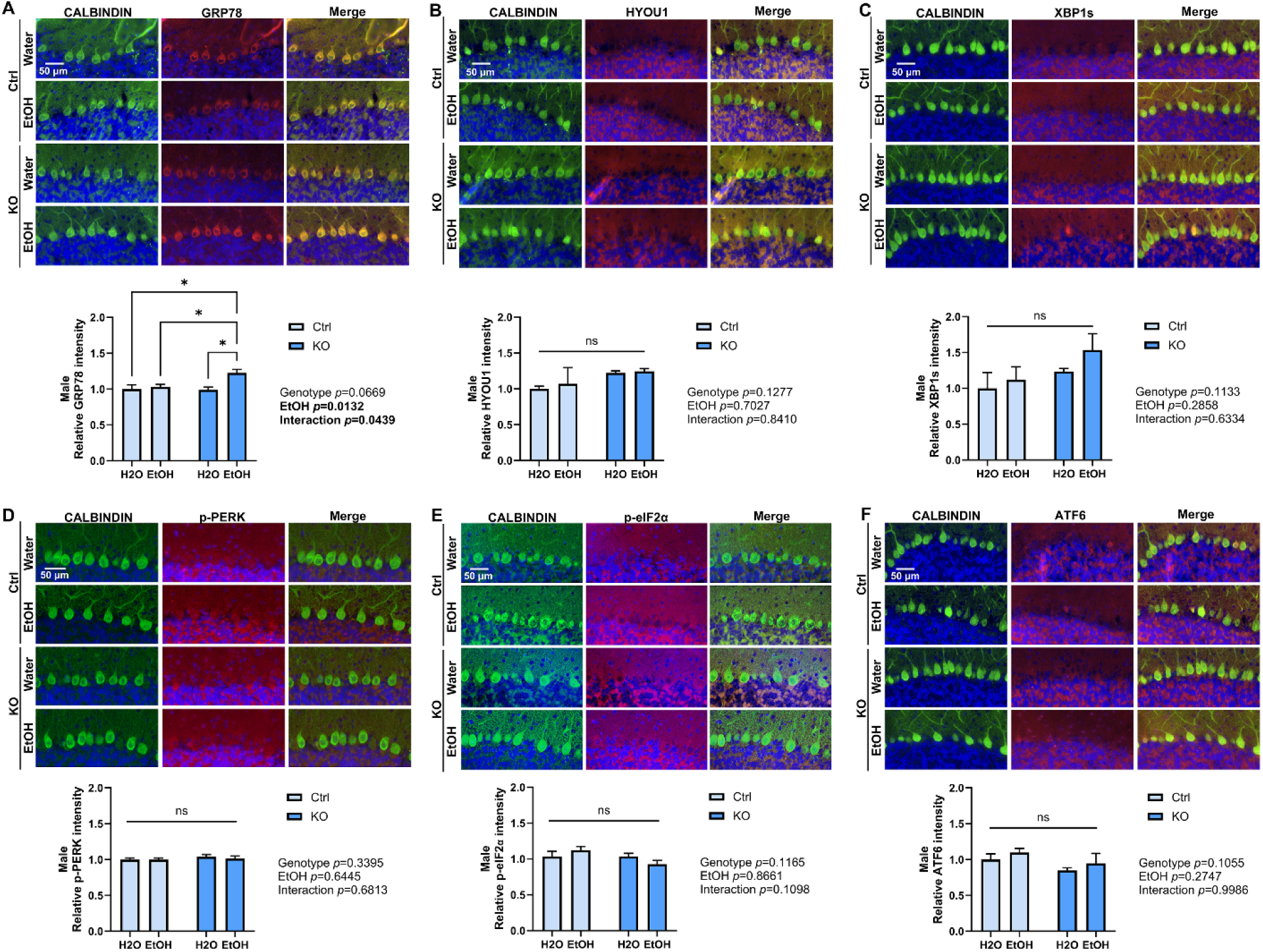
Effects of ethanol exposure on ER stress in male MANF KO PCs. **A-F.** Representative immunofluorescent images and quantifications for the expression of ER stress markers GRP78 (A), HYOU1 (B), XBP1s (C), p-PERK (D), p-eIF2α (E), and ATF6 (F) in water- and ethanol-exposed control and KO male mice cerebellum. PC marker Calbindin was co-labeled in green. Quantification of ER stress markers fluorescence intensity was expressed as mean ± SEM. n=3-4 per group. Two-way ANOVA followed by Tukey’s *post hoc* test. Significant main effect *p* values were highlighted in bold; \**p*< 0.05; ns not significant.

To further determine whether alcohol and MANF KO result in PC cell death and degeneration, cerebellum vermis sections were immunolabeled with apoptotic marker cleaved caspase-3 and tested in TUNEL assay. In line with alcohol and MANF KO induced female specific PCs shrinkage and UPR activation, we observed a significant increase in the number of cleaved caspase-3 positive and TUNEL positive PCs in ethanol treated female MANF KO cerebellum (Figure 8A and 8C), while no difference in PCs apoptosis was detected in males (Figure 8B and 8D).

**Figure 8.**
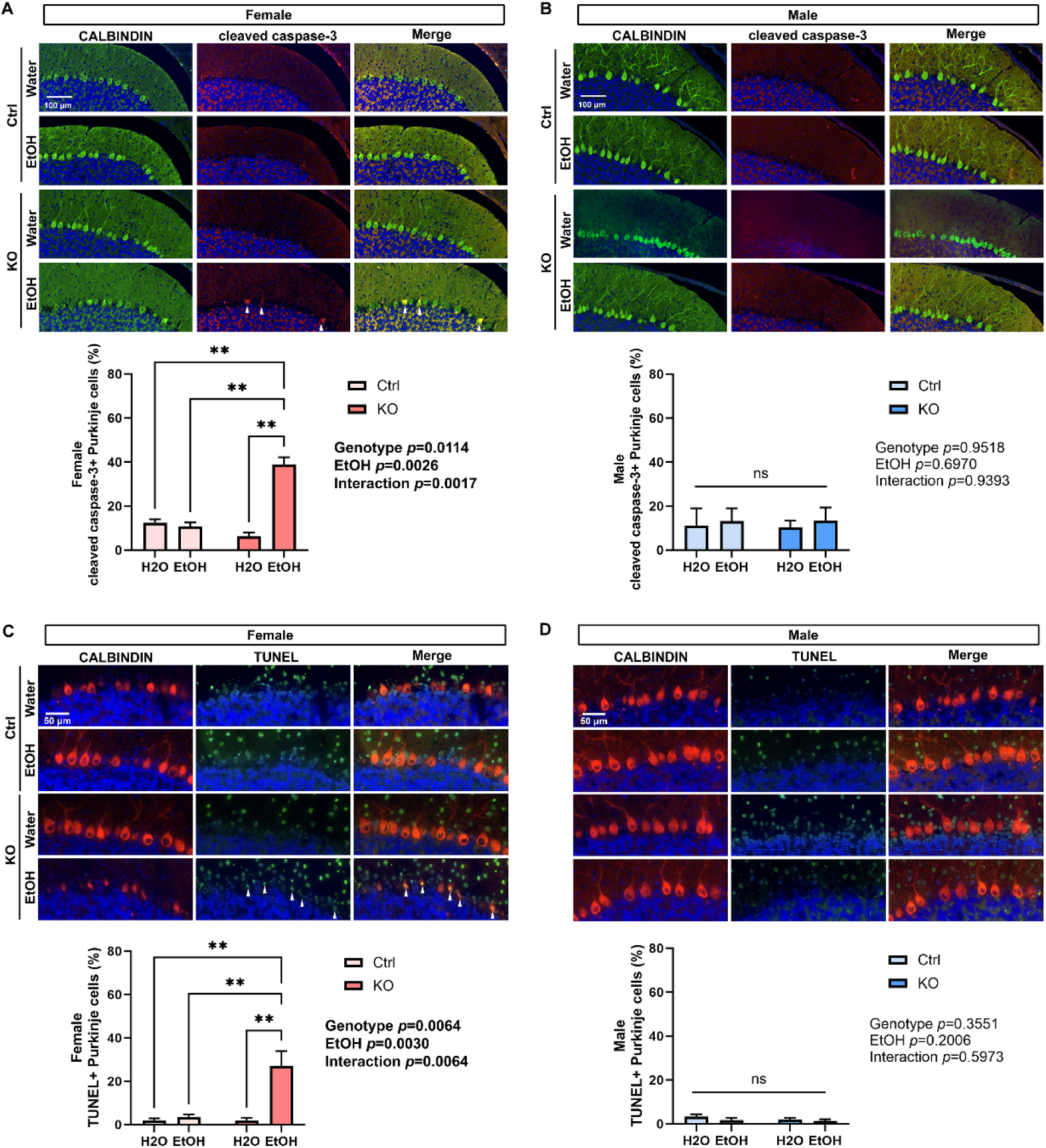
Ethanol induces apoptosis in female but not male MANF KO PCs. **A-B.** Representative immunofluorescent images of cleaved caspase-3 (red) and PC marker Calbindin (green) in water- and ethanol-exposed control and KO female (A) and male (B) cerebellum. The number of cleaved caspase-3 positive PCs in lobule II was quantified and expressed as mean ± SEM. n=3-4 per group. Two-way ANOVA followed by Tukey’s *post hoc* test. Significant main effect *p* values were highlighted in bold; \*\**p*< 0.01; ns not significant. **C-D.** Representative images of TUNEL labeling (green) with immunofluorescent co-labeling of Calbindin (red) in water- and ethanol-exposed control and KO female (C) and male (D) cerebellum. The number of TUNEL positive apoptotic PCs in lobule II was quantified and expressed as mean ± SEM. n=3-4 per group. Two-way ANOVA followed by Tukey’s *post hoc* test. Significant main effect *p* values were highlighted in bold; \*\**p*< 0.01; ns not significant. Arrow heads in A and C indicate cleaved caspase-3 and TUNEL positive PCs, respectively.

### MANF deficiency altered the transcriptomic signature in PCs

To comprehensively understand how MANF deficiency changes the transcriptomic landscape in PCs, and how it contributes to the female susceptibility in alcohol neurotoxicity, we used Visium spatial transcriptomics analysis (STA) to profile gene expression in the cerebellum vermis from age matched 4 months old mice, including 3 control males, 3 KO males, 3 control females, and 3 KO females. Samples were arranged on 3 Visium slides as illustrated in Figure 9A. PCs specific MANF deficiency was confirmed by MANF immunostaining for all the KO samples before the analysis (Figure S6A). We detected 1,185-3,492 Visium spots per sample and 4,478-6,641 genes per spot, which lead to more than 20,000 genes that were sequenced per sample.

**Figure 9.**
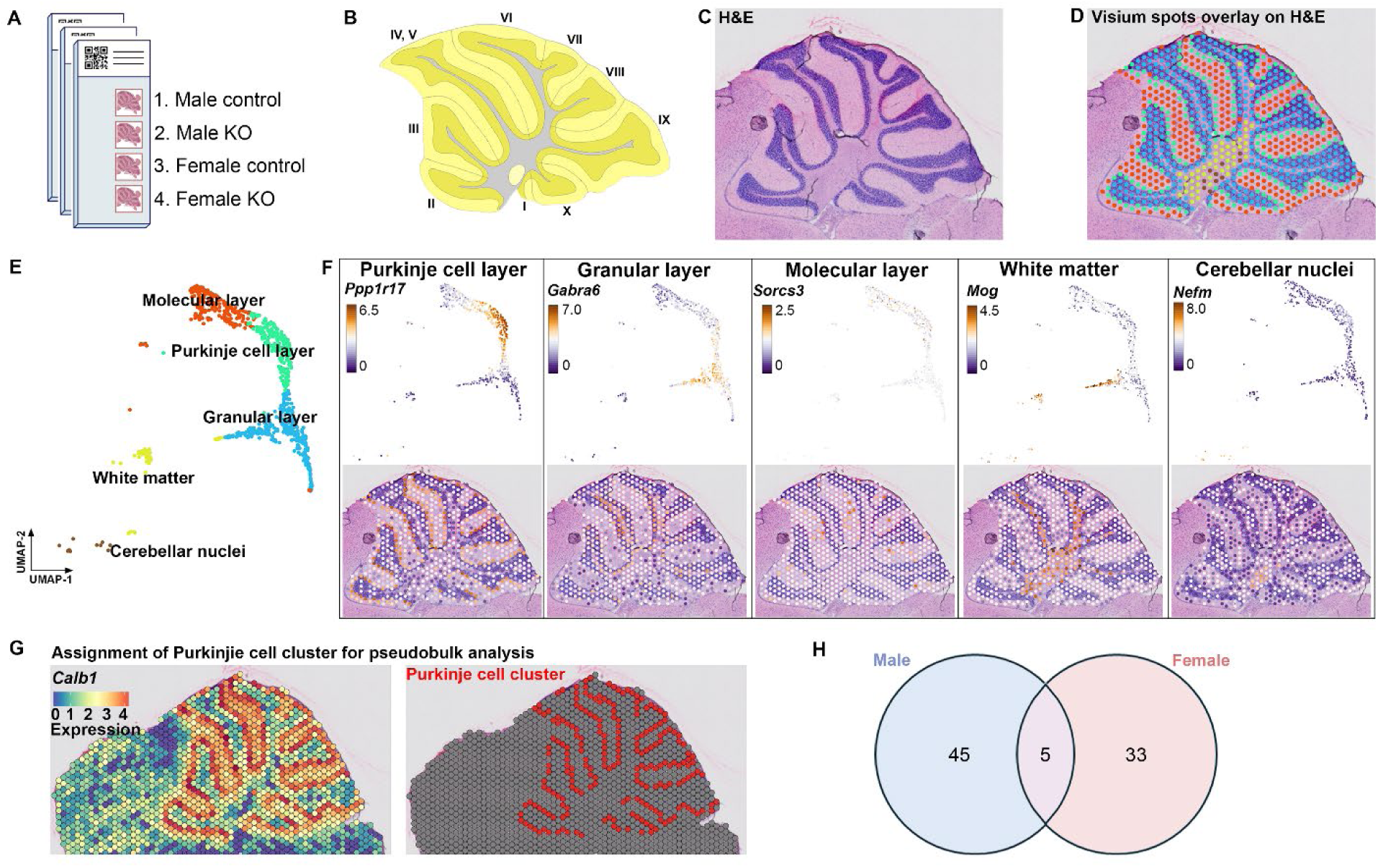
Visium spatial transcriptomics analysis identifies spatially defined tissue clusters in the mouse cerebellum. **A.** Layout of the 3 Visium slides with sagittal sections of the cerebellum vermis from male and female control and MANF KO adult mice. **B.** Mouse cerebellum vermis sagittal section (adapted screenshot from the Allen Reference Atlas, image 17 of 21, position 56). Cerebellar lobules were indicated by Roman numerals. **C.** Example of H&E staining image of Visium slide 1, sample 1. **D.** H&E staining image overlayed with unsupervised graph-based clustering of Visium spots in the cerebellum (spots outside of the cerebellum were not shown). **E.** UMAP plot of the Visium spots with annotated tissue clusters. **F.** Expression of selected marker genes for each tissue cluster in the cerebellum. **G.** PC cluster was selected for further pseudobulk analysis based on *Calbindin* (*Calb1*) transcripts distribution in the cerebellum. **H.** Venn diagram showing the number of male and female overlapping and distinct differentially expressed genes (DEGs) in the PC cluster.

Unsupervised clustering identified spots with defined tissue clusters and marker genes expression that recapitulated the cerebellum vermis regions, including the Purkinje cell layer, granular layer, molecular layer, white matter, and cerebellar nuclei (Figure 9B-9F). Spots for PC cluster were selected based on *Calbindin* (*Calb1*) transcripts distribution in the cerebellum, which were extracted for further pseudobulk analysis (Figure 9G). PC cluster from female controls were compared with that of female KOs to identify differentially expressed genes (DEGs) in female, and male controls were compared with that of male KOs to identify DEGs in male. We identified 38 DEGs in female and 50 DEGs in male, among which 5 were shared in both female and male, 33 were female specific, and 45 were male specific (Figure 9H, Table S1). Enrichment analysis indicated that many of the DEGs were involved in protein binding and amino acid transmembrane transportation (Figure 10-12). Considering that the Visium spots did not provide a single cell resolution, thus the results may be interfered by genes from cells surrounding the PCs such as the Bergmann glia, granular cell, and molecular layer neurons, we further employed Xenium in situ analysis with high resolution single cell information to validate the Visium results and to find new genes that may be potentially overlooked in Visium. A customized 300-gene panel was designed that included most of the DEGs identified in Visium that met the Xenium probe design criteria (Table S2). In addition, it also contained genes encoding MANF interacting proteins reported in previous publications and in the STRING database (version 12.0) [31, 72], genes involved in ER stress/UPR, neuroinflammation, and oxidative stress, and cerebellum cell type marker genes [73] (Table S2). Cerebellum vermis sections from age matched 4 months old mice, including 3 control males, 3 KO males, 3 control females, and 3 KO females, were confirmed with PC specific MANF KO by immunostaining before proceeding to Xenium (Figure S6B). We detected 52,901-75,418 cells per sample with 100-132 transcripts per cell. Cell segmentation boundaries were visualized and transcripts of the 300 genes were assigned to each cell in Xenium Explorer (Figure 13A). Segregated cells were classified into different cell types using unsupervised graph-based clustering and the distribution of cell type marker genes were consistent with the cell clusters (Figure 13B). Cells in the PC cluster were selected based on the PC specific gene *Ppp1r17* (protein phosphatase 1 regulatory subunit 17) transcripts expression and were extracted for further pseudobulk analysis (Figure 13C). We identified 14 DEGs in female PCs and 16 DEGs in male PCs (Figure 14A, 14B; Table S3). Among the DEGs, 10 of them were shared by male and female, and all 10 common DEGs were upregulated in KO PCs (Figure 15A). Enrichment analysis indicated that most of these common DEGs, including *Hspa5*, *Hsp90b1*, *Hyou1*, *P4hb*, *Pdia6*, *Xbp1*, and *Sdf2l1*, were involved in protein folding and response to ER stress (Figure 15B). In addition, it should be noted that the upregulation of two DEGs, *Xab2* and *Camsap3*, were potentially not due to MANF KO, but because of the cloning of the mouse genome sequence of 3,436,600-3,609,640 on chromosome 8 flanking *Pcp2* gene into the bacterial artificial chromosome (BAC) while generating the *Pcp2-Cre* line [63]. The intact *Xab2* and *Camsap3* genomic sequence was contained in this genome region, thus potentially resulted in their upregulation in KO. Other than the 10 DEGs shared by female and male, four additional DEGs were found to be female-specific, among which three were ER related, including *Creld2*, *Ddost*, and *Pdia3.* They showed female specific upregulation in MANF KO PCs. *Creld2* had protein disulfide isomerase activity, while *Ddost* and *Pdia3* were in ER protein-containing complex and function in protein processing in ER (Figure 15C and 15D). *Kat2b* was downregulated in female KO PCs and was involved in cellular response to nutrient (Figure 15C and 15D). Six additional DEGs were male-specific. *Camk2n1* and *Derl3* were upregulated in male MANF KO PCs, while *Fos*, *Gabra6*, *Rbfox3* and *Txnip* were downregulated with *Fos* and *Txnip* involved in response to progesterone (Figure 15E and 15F). When comparing the Visium and Xenium results, the upregulation of *Cfap100* in MANF KO PCs was confirmed in both Visium and Xenium analyses, although it showed up only in female in Visium but in both female and male in Xenium. The downregulation of *Fos* and *Txnip* in male MANF KO PCs were also confirmed in both Visium and Xenium.

**Figure 10.**
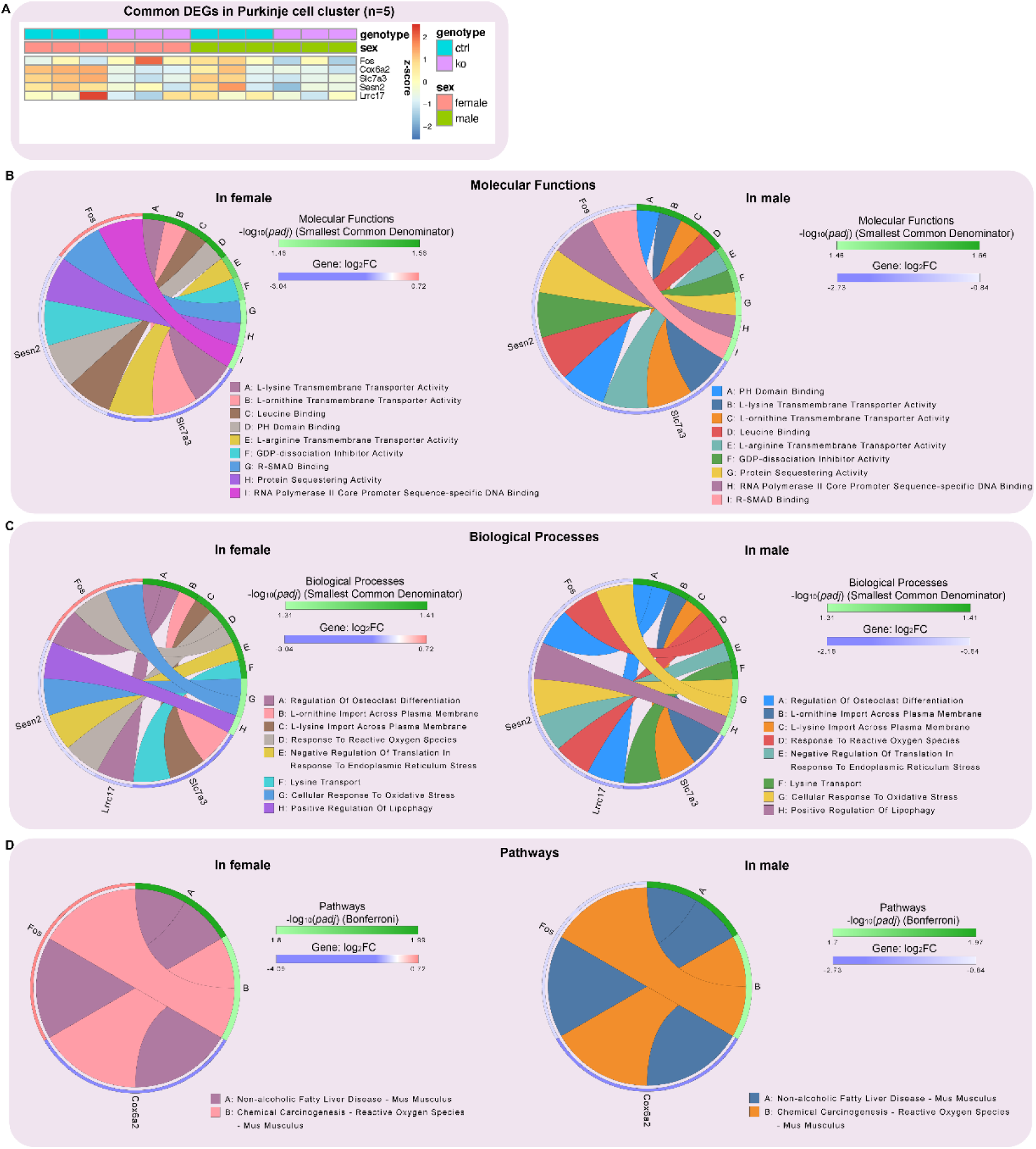
Transcriptional impact of MANF deficiency in both female and male PCs. **A.** Heatmap of differentially expressed genes (DEGs) in control and MANF KO PCs shared in both female and male PC cluster identified by pseudobulk analysis. False discovery rate (FDR)-adjusted *p* value< 0.1, |log2 FC|> 0.2, and baseMean value> 5. **B.** Common DEGs enrichment analysis using the Gene Ontology (GO) consortium database for molecular functions. The circular plot shows the top molecular functions and their most important genes in the PCs that are affected by MANF KO. Smallest common denominator-adjusted *p* value<0.05, that is -log10 (*padj*)>1.3. **C.** Common DEGs enrichment analysis using the GO dataset for biological processes. The circular plot shows the top biological processes and their most important genes in the PCs that are affected by MANF KO. Smallest common denominator-adjusted *p* value<0.05, that is -log10 (*padj*)>1.3. **D.** Common DEGs enrichment analysis using the Kyoto Encyclopedia of Genes and Genomes (KEGG) database for pathways. The circular plot shows the top pathways and their most important genes in the PCs that are affected by MANF KO. Bonferroni-adjusted *p* value<0.05, that is -log10 (*padj*)>1.3. Upregulated genes are shown in red, downregulated genes are shown in blue.

**Figure 11.**
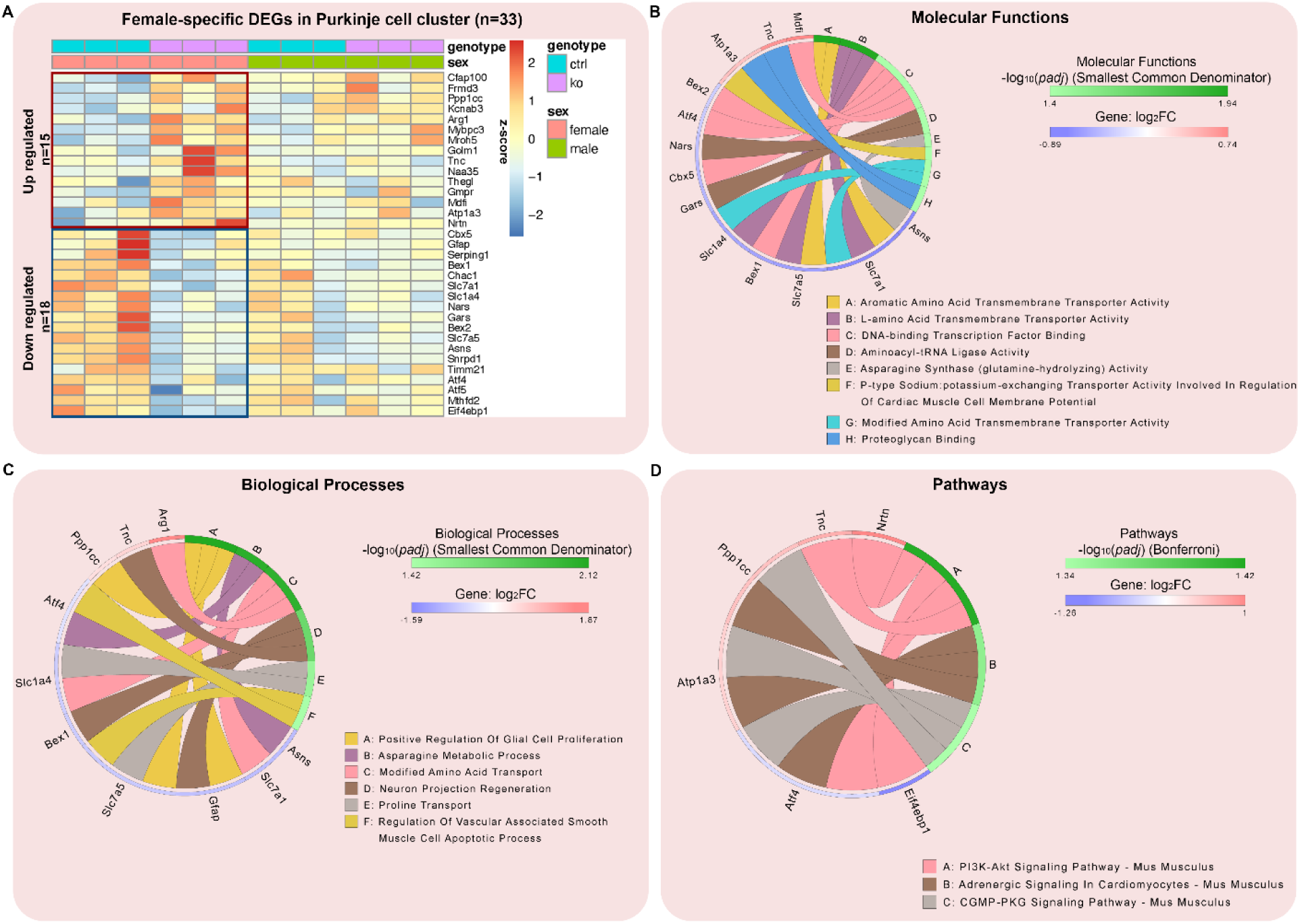
Female specific transcriptional changes in PCs in response to MANF deficiency. **A.** Heatmap of female specific DEGs in control and MANF KO PCs identified by PC cluster pseudobulk analysis. FDR-adjusted *p* value<0.1 and |log2 FC|> 0.2. **B.** Female specific DEGs enrichment analysis using the Gene Ontology (GO) consortium database for molecular functions. The circular plot shows the female specific top molecular functions and their most important genes in the PCs that are affected by MANF KO. Smallest common denominator-adjusted *p* value<0.05, that is -log10 (*padj*)>1.3. **C.** Female specific DEGs enrichment analysis using the GO consortium database for biological processes. The circular plot shows the female specific top biological processes and their most important genes in the PCs that are affected by MANF KO. Smallest common denominator-adjusted *p* value<0.05, that is -log10 (*padj*)>1.3. **D.** Female specific DEGs enrichment analysis using the Kyoto Encyclopedia of Genes and Genomes (KEGG) database for pathways. The circular plot shows the female specific top pathways and their most important genes in the PCs that are affected by MANF KO. Bonferroni-adjusted *p* value<0.05, that is -log10 (*padj*)>1.3. Upregulated genes are shown in red, downregulated genes are shown in blue.

**Figure 12.**
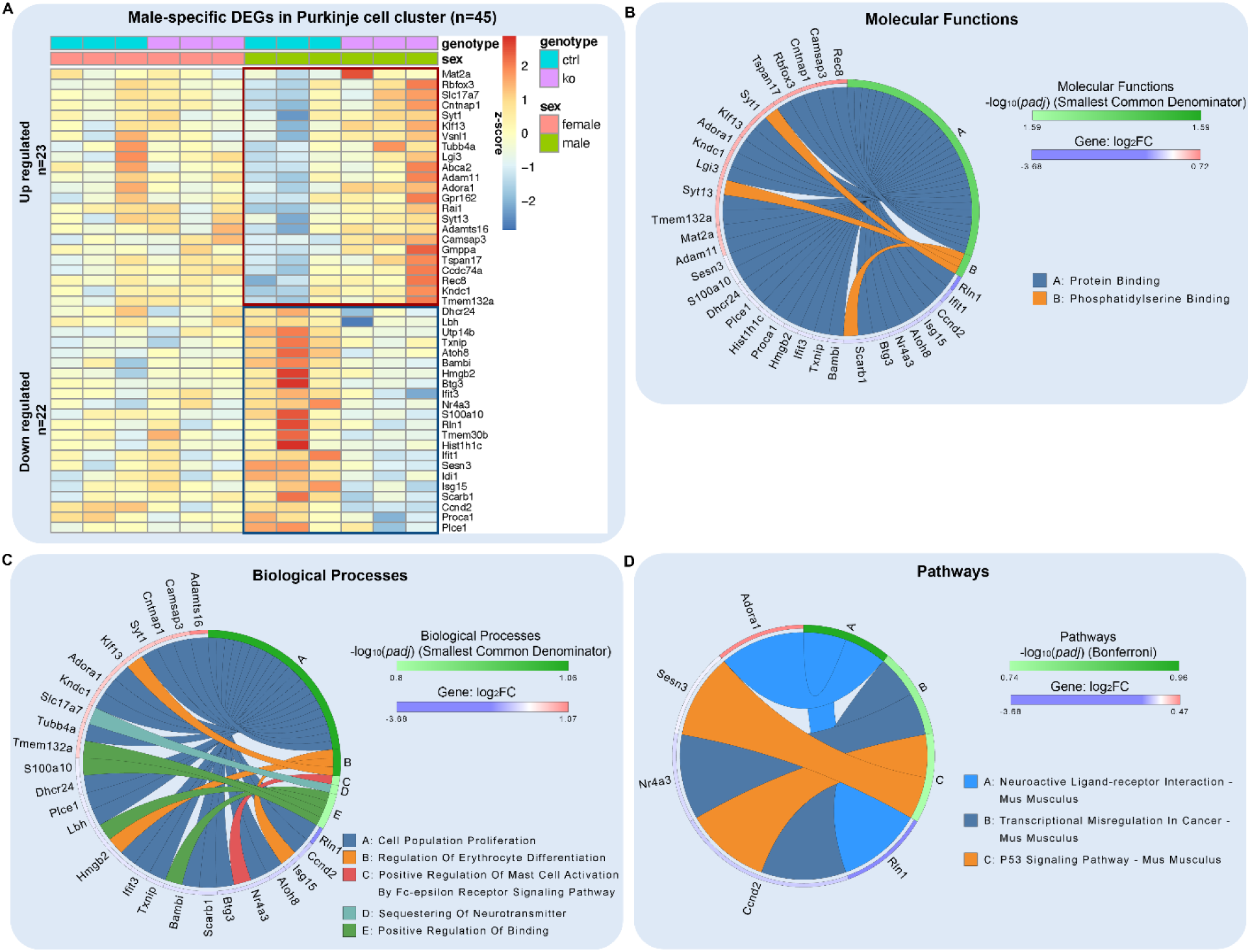
Male specific transcriptional changes in PCs in response to MANF deficiency. **A.** Heatmap of male specific DEGs in control and MANF KO PCs identified by PC cluster pseudobulk analysis. FDR-adjusted *p* value<0.1 and |log2 FC|> 0.2. **B.** Male specific DEGs enrichment analysis using the Gene Ontology (GO) consortium database for molecular functions. The circular plot shows the male specific top molecular functions and their most important genes in the PCs that are affected by MANF KO. Smallest common denominator-adjusted *p* value<0.05, that is -log10 (*padj*)>1.3. **C.** Male specific DEGs enrichment analysis using the GO consortium database for biological processes. The circular plot shows the male specific top biological processes and their most important genes in the PCs that are affected by MANF KO. No processes were identified with the significant smallest common denominator-adjusted *p* values<0.05. The top 5 processes with the smallest adjusted *p* values were listed. **D.** Male specific DEGs enrichment analysis using the Kyoto Encyclopedia of Genes and Genomes (KEGG) database for pathways. The circular plot shows the male specific top pathways and their most important genes in the PCs that are affected by MANF KO. No pathways were identified with the significant Bonferroni-adjusted *p* value<0.05. The top 3 pathways with the smallest adjusted *p* values were listed. Upregulated genes are shown in red, downregulated genes are shown in blue.

**Figure 13.**
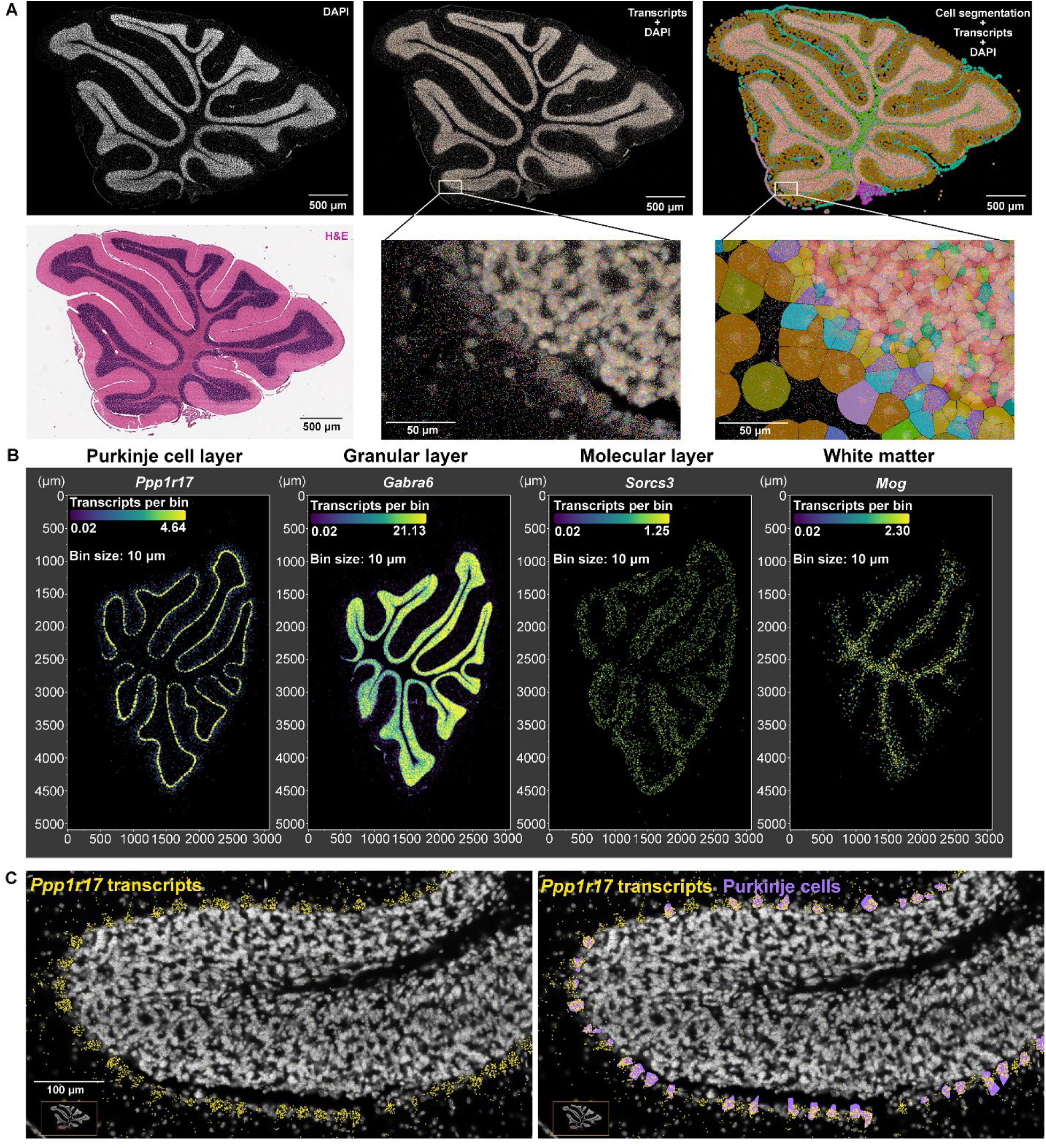
Xenium-based single cell in situ gene expression profile in control and MANF KO cerebellum. **A.** Example of Xenium slide 1, sample 1 cerebellum vermis section stained with DAPI and H&E. Transcripts for 300 genes and unsupervised graph-based clustering of segregated cells were overlayed with DAPI image. Zoom-in images demonstrated individual transcripts as colored dots and cells filled with colors representing different clusters. **B.** Transcript density maps of selected marker genes for distinct cell clusters in the cerebellum. **C.** Cells in the PC cluster (purple) were selected for further pseudobulk analysis based on *Ppp1r17* transcripts expression (yellow dots) in the cerebellum.

**Figure 14.**
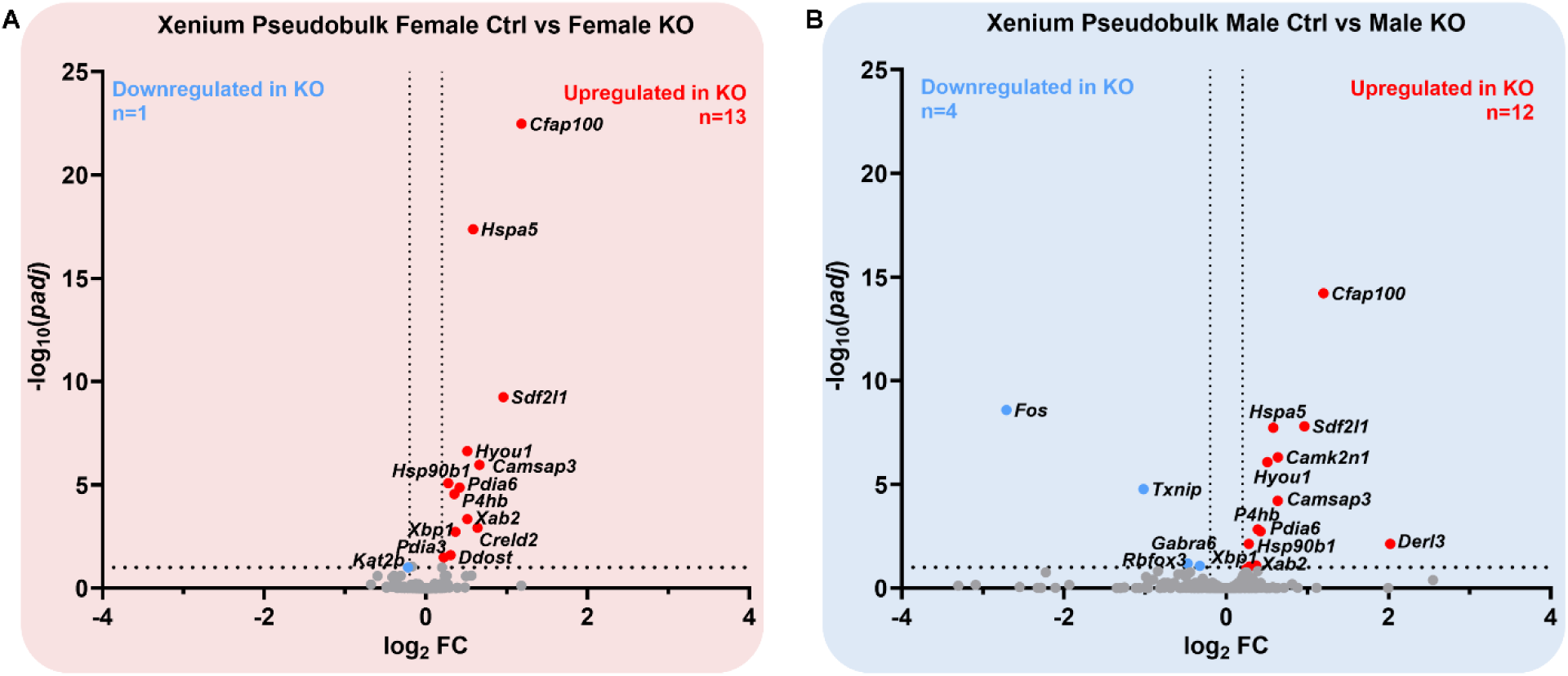
Xenium pseudobulk analysis of PC cluster identifies DEGs in female and male in response to MANF deficiency. **A-B.** Vulcano plot showing the -log10 (*padj*) and log2 FC for the 300 genes in the Xenium gene panel in female (A) and male (B). FC: fold change. Black dotted lines indicate the threshold of -log10 (*padj*)>1, that equaled to FDR-adjusted *p* value<0.1, and |log2 FC|> 0.2. Red dots: upregulated genes in MANF KO PCs; blue dots: downregulated genes in MANF KO PCs; grey dots: not significantly changed genes.

**Figure 15.**
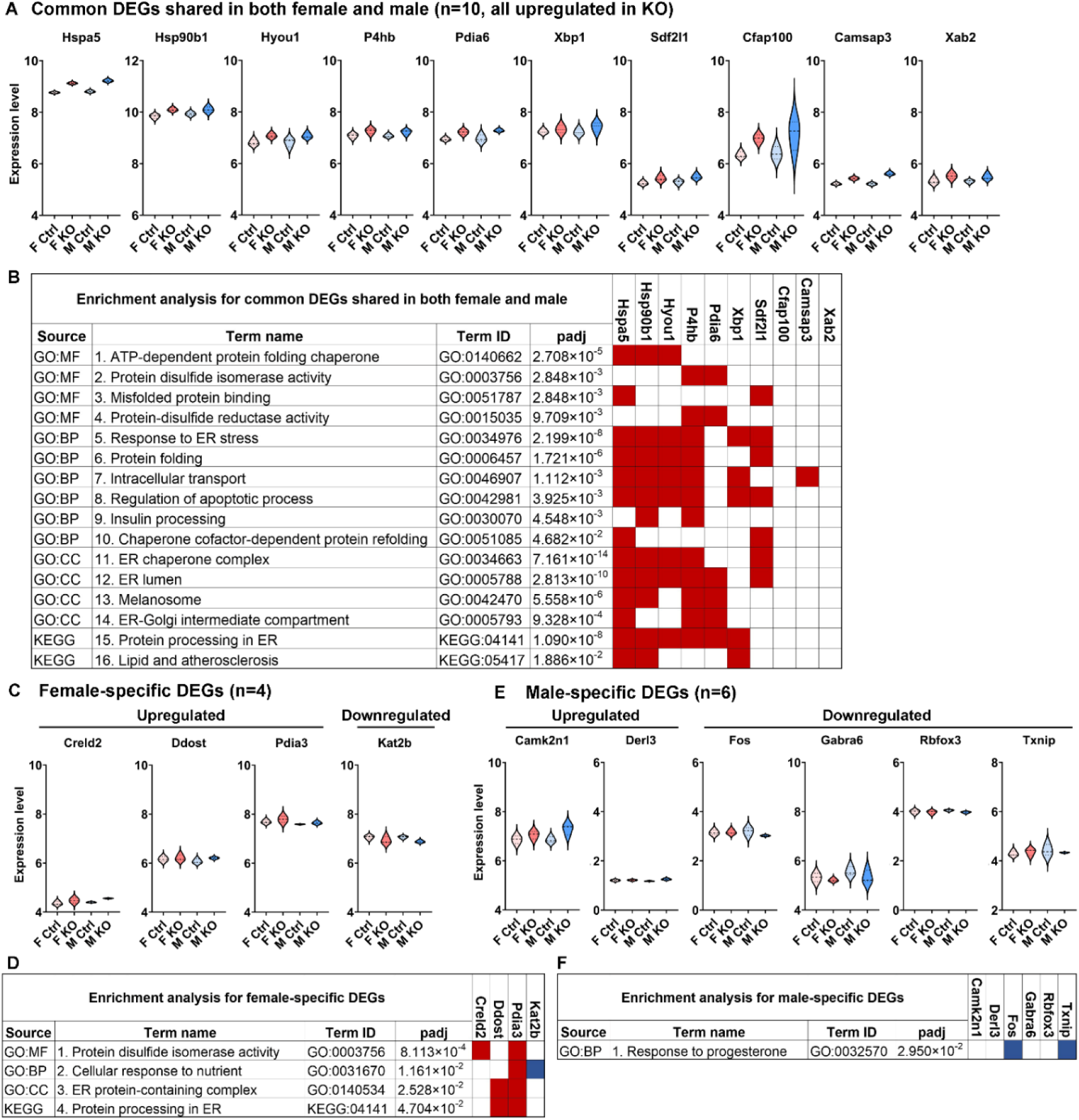
Enrichment analysis of DEGs identified in Xenium PC pseudobulk analysis indicates upregulation of ER related genes in MANF KO PCs. **A.** Violin plots for the expression levels of common DEGs shared in both female and male control and MANF KO PCs. **B.** Enrichment analysis for the common DEGs. **C.** Violin plots for the expression levels of female-specific DEGs in control and MANF KO PCs. **D.** Enrichment analysis for the female-specific DEGs. **E.** Violin plots for the expression levels of male-specific DEGs in control and MANF KO PCs. **F.** Enrichment analysis for the male-specific DEGs. Data sources used in the analysis include Gene Ontology (GO) consortium database and Kyoto Encyclopedia of Genes and Genomes (KEGG) database. MF: molecular function; BP: biological process; CC: cellular component. Adjusted *p* value (padj) was calculated using g:SCS (Set Counts and Sizes) method developed by g:Profiler. Driver GO terms and KEGG terms with padj<0.05 were listed. The table demonstrated the source of database, term names and IDs with genes involved in each term. Red square: the gene was involved in the term and was upregulated; blue square: the gene was involved in the term and was downregulated.

## Discussion

In this study, we generated cerebellar PC specific MANF KO mouse model and investigated the effect of MANF deficiency on alcohol induced behavioral impairments and cellular and molecular changes in PCs in the adult mice. We also identified target genes that their expression was altered in response to the loss of MANF, potentially contributing to the phenotype. We found that adult mice with PC specific MANF deficiency exhibit impaired motor functions. Binge alcohol exposure interacted with MANF deficiency to exacerbate the motor deficits in these animals. Interestingly, female KOs were more sensitive than male KOs to alcohol induced motor function impairments. In line with the behavior results, alcohol treatment led to UPR activation, alteration in the intracellular distribution of calcium binding protein Calbindin, and PC degeneration in female MANF deficient PCs but not males. Spatial transcriptomics and high throughput in situ analyses demonstrated that MANF deficiency led to sex specific alterations in the transcriptomic landscape in PCs and triggered the expression of genes involved in protein folding and response to ER stress, potentially contributing to alcohol induced cellular damages in MANF KO PCs. These results suggests that MANF deficient PCs may be predisposed with a higher risk to UPR activation and ER stress in a sex dependent manner, contributing to their vulnerability to alcohol neurotoxicity.

This study highlighted the importance of protein homeostasis in PCs survival and alcohol neurotoxicity. PCs have a high demand for protein synthesis and are vulnerable to disruptions in ER function. Conditional knockout of the major ER chaperone GRP78 in PCs resulted in UPR activation and PC degeneration [74]. Imbalance in protein homeostasis is associated with PC degeneration in various disease models. For example, upregulation of UPR was observed in the woozy (wz) mutant mouse that exhibit cerebellar ataxia and PC loss due to mutation in the GRP78 nucleotide exchange factor gene *Sil1*, which was associated with the Marinesco-Sjögren syndrome in human [75], demonstrating that perturbation of ER chaperone function in PCs cause ER stress and subsequent PC degeneration [76, 77]. Similarly, in the Purkinje cell degeneration (pcd) transgenic mouse model that exhibit progressive PC degeneration and ataxia, a significant increase in the expression of GRP78 was observed, along with the upregulation of ER stress induced apoptosis markers CHOP and caspase 12, suggesting that the activation of UPR contributed to PC degeneration in pcd mouse model [78]. ER stress is also associated with PC degeneration in several mouse models of SCA, including SCA type 6, 17, and 22 [79–81]. In addition, ER-associated degradation (ERAD) is a conserved quality control pathway responsible for the removal of misfolded proteins in the ER. PC-specific deletion of Sel1L (suppressor of lin-12-like 1), which is a key ERAD adaptor protein, led to progressive PC loss in mice [82]. Previously, we have shown that alcohol caused widespread ER stress and UPR activation in the developing and adult mouse brain [26, 28]. Chronic alcohol exposure has been reported to increase the expression of ER stress markers and induce dilation of the smooth ER in PCs in adult and aging rat brain [7–9]. Given the sensitivity of PCs to ER stress, alcohol induced ER dysfunction and imbalance in protein homeostasis may contribute to the vulnerability of PCs to the toxic effect of alcohol. In this study, binge alcohol exposure or MANF deficiency separately did not induce PC degeneration. However, when combine alcohol with MANF deficiency, significant PC degeneration was observed, especially in females. This may be largely due to the function of MANF in modulating ER stress.

MANF expression and secretion can be induced by ER stressors, indicating that MANF is involved in ER stress related signaling pathways. Various *in vitro* and *in vivo* studies have demonstrated that under ER stressed conditions such as cerebral ischemia, hypoxia, and some neurodegenerative diseases, MANF deficiency led to elevated UPR and cell death, while MANF overexpression promoted neuronal survival [47, 51, 55, 83, 84]. Previously using a neuron-specific MANF KO mouse model, we found that MANF KO mice were compromised in ER homeostasis modulation and were more susceptible to alcohol-induced neuronal ER stress and neurodegeneration in the developing and mature mouse brain [31, 32, 85]. The current study supported these finding and further demonstrated that MANF deficiency and alcohol exposure synergistically induced ER stress and impaired PCs function. The exact molecular mechanism of how MANF modulates ER stress is poorly understood. One possible mechanism is through its physical interaction with the key ER chaperone GRP78. MANF was found to bind to the adenosine diphosphate (ADP)-bound GRP78 to inhibit ADP release from GRP78 and stabilize the complex of GRP78 with its substrate proteins, facilitating protein folding [45, 53]. MANF also form protein interactome with several additional proteins that were involved in protein folding, including a GRP78 nucleotide exchange factor HYOU1 and protein disulfide isomerases PDIA1 and PDIA6 [72]. Genes encoding these MANF interacting proteins, including *Hspa5* (encodes for GRP78), *Hyou1*, *P4hb* (encodes for PDIA1), and *Pdia6* were all identified to be upregulated in MANF KO PCs in our Xenium analysis. In addition, we also found several more ER stress related genes that were upregulated in MANF KO PCs, including *Hsp90b1*, *Xbp1*, and *Sdf2l1*. These results suggest that a compensatory transcriptional activity occurred in MANF deficient PCs, although fully compensation is unlikely as when treated with alcohol, MANF KO PCs still displayed elevated UPR and neurodegeneration comparing to controls. The cytoprotective role of MANF was also suggested to be independent to its binding with GRP78. Eesmaa et al reported that MANF mutant proteins that were unable to bind to GRP78 can still promote survival of primary mouse superior cervical ganglion (SCG) neuron cultures in vitro [72]. In fact, recently the same group found that MANF can bind to and negatively regulate the ER transmembrane UPR sensor protein IRE1α [86]. MANF-IRE1α interaction attenuated UPR and was required for the pro-survival activity of MANF. TXNIP is a IRE1α downstream target gene as a key regulator in cellular stress. It can induce oxidative stress, inflammation and apoptosis [87]. Interestingly, our Visium and Xenium results both indicated a downregulation of *Txnip* in male MANF KO PCs but not female, suggesting that male and female PCs may have distinct molecular responses to MANF deficiency and hence a sex-specific sensitivity to alcohol neurotoxicity. Additional sex-specific ER related genes were identified in our Xenium results, include *Ddost*, *Creld2*, and *Pdia3*, which were upregulated in female but not in male MANF KO PCs.

It is not surprising to find sex specific response to alcohol in MANF deficient PCs. Sex dimorphism in the effect of alcohol have been reported in human and animal studies that females were more vulnerable than males to alcohol induced organ damages, including the brain [88, 89]. Although women tend to drink less volume and shorter periods of alcohol than men, women exhibit greater volumetric brain loss and increased cognitive impairment than men [90, 91]. The etiology of this sex difference in alcohol neurotoxicity remains poorly understood. Differences in alcohol metabolism, brain structure, hormone, and genetics between male and female may all contribute to the female vulnerability. Research in animal models have demonstrated that for the same amount and period of alcohol exposure, females showed greater neuronal loss [92], severer neuroinflammation [93, 94], different neuroadaptive responses to alcohol withdrawal [95], and distinct transcriptomics alterations [96] than males. We have shown previously using neuronal MANF deficient adult mice that female, but not male MANF KO mice exhibited an increased locomotor activity and female MANF KO animals were more susceptible to alcohol induced body weight loss [32]. We also found that alcohol altered the neuronal expression of several UPR and neuroinflammation markers in MANF KO mice in a sex-specific manner [32]. The present findings provide additional evidence to support the sex-specific effect of MANF deficiency on alcohol neurotoxicity, emphasizing the importance to consider sex difference in alcohol research.

Another interesting phenotype we observed was the intranuclear translocation of Calbindin in alcohol treated female MANF KO PCs. Calbindin is a calcium binding protein acts primarily as a cellular Ca^2+^ buffer in PCs. Calbindin can regulate neuronal calcium homeostasis and modify calcium-dependent intracellular signaling by buffering calcium and modulating calcium channel activity [97]. Calbindin protein normally distributes diffusely in the cytoplasm, dendrites, axon, and nucleus of PCs. Increased PCs intranuclear Calbindin expression has been reported in animal models with hypoxia and chronic morphine treatment [98, 99], indicating an increased intranuclear Ca^2+^ concentration, that may regulate gene transcription and activate intranuclear Ca^2+^ signaling in PCs [100]. The ER is the primary intracellular Ca^2+^ storage organelle in eukaryotic cells. The ER surrounding the nucleus is contiguous with the outer nuclear membrane of the nuclear envelope, referred to as the nucleoplasmic reticulum (NR). Ca^2+^ can enter the nucleus from the cytoplasm through nuclear pores or can be released from the NR via specific channels, such as the ligand-gated Ca^2+^ release channels inositol 1,4,5-trisphosphate receptor (InsP3R) and ryanodine receptors (RyRs) [101, 102]. The dynamic structure of the NR can be altered in pathological conditions [103]. Disruption of Ca^2+^ homeostasis and increased intracellular Ca^2+^ is a key mechanism underlying alcohol-induced neuronal death. Long term alcohol exposure was reported to induce InsP3R upregulation in mouse cerebral cortical neurons [104]. Study using primary cultures of cerebellar granule neurons has demonstrated that alcohol can induce increased intracellular Ca^2+^ and pretreatment with InsP3R inhibitor blocked both alcohol-induced rise in Ca^2+^ and neuronal death [105]. Alcohol induced increase in intranuclear Calbindin expression in female MANF KO PCs suggests that MANF may be involved in regulating the cellular response to alcohol-induced Ca^2+^ dysregulation. MANF deficiency may potentially alter the intracellular Ca^2+^ signaling in response to alcohol and contribute to alcohol neurotoxicity.

In conclusion, the current findings demonstrate that MANF deficient PCs are more susceptible to binge alcohol exposure induced ER stress and neurodegeneration in the adult brain in a sex dependent manner. Such vulnerability is particularly increased in MANF deficient female PCs. We identified overlapping and sex-specific differentially expressed genes in the adult PCs, indicating an alteration in the PC transcriptomics landscape in response to MANF deficiency. Many of the altered genes are involved in protein binding, amino acid transmembrane transportation, and facilitating protein folding. Our study has established a valuable animal model to investigate the role of MANF in PC structure and function as well as molecular mechanisms underlying alcohol-induced PC damages. Further studies are to focus on the biological function of the altered genes, and how they contribute to the increased susceptibility of MANF deficient neurons to alcohol neurotoxicity in the female brain.

## Funding

This work was supported by the National Institute on Alcohol Abuse and Alcoholism (AA017226 and AA015407).

## Acknowledgement

This work was supported by the National Institutes of Health (NIH) grants AA017226 and AA015407. We thank Marisol Lauffer for performing and analyze data from the Barnes maze test and we acknowledge the personnel and instrumentation in the Neural Circuits and Behavior Core in the Iowa Neuroscience Institute, supported in part by the Roy J. Carver Charitable Trust and UI Carver College of Medicine. We thank the Comparative Pathology Laboratory and the Iowa Neuropathology Resource Laboratory at University of Iowa Department of Pathology for providing histology services. Visium sample preparation, sequencing, and data analysis presented herein were obtained at the Iowa NeuroBank Core in the Iowa Neuroscience Institute, and the Genomics Division and Bioinformatics Division of the Iowa Institute of Human Genetics, which is supported, in part, by the University of Iowa Carver College of Medicine and the Holden Comprehensive Cancer Center (National Cancer Institute of the National Institutes of Health under Award Number P30CA086862). We thank the University of Michigan Advanced Genomics core for providing Xenium in situ service supported by the National Cancer Institutes of Health (P30CA046592) using single cell and spatial analysis as Cancer Center Shared Resource. We thank the Genomics Research and Technology Hub (GRT Hub), University of California Irvine (UCI) and Dr. Jie (Jenny) Wu for performing Xenium data analysis that utilized resources of the UCI GRT Hub parts of which are supported by NIH grants to the Comprehensive Cancer Center (P30CA062203) and the UCI Skin Biology Resource Based Center (P30AR075047) at UCI, as well as to the GRT Hub for instrumentation (1S10OD010794-01and 1S10OD021718-01).

---

**Figure S1.**
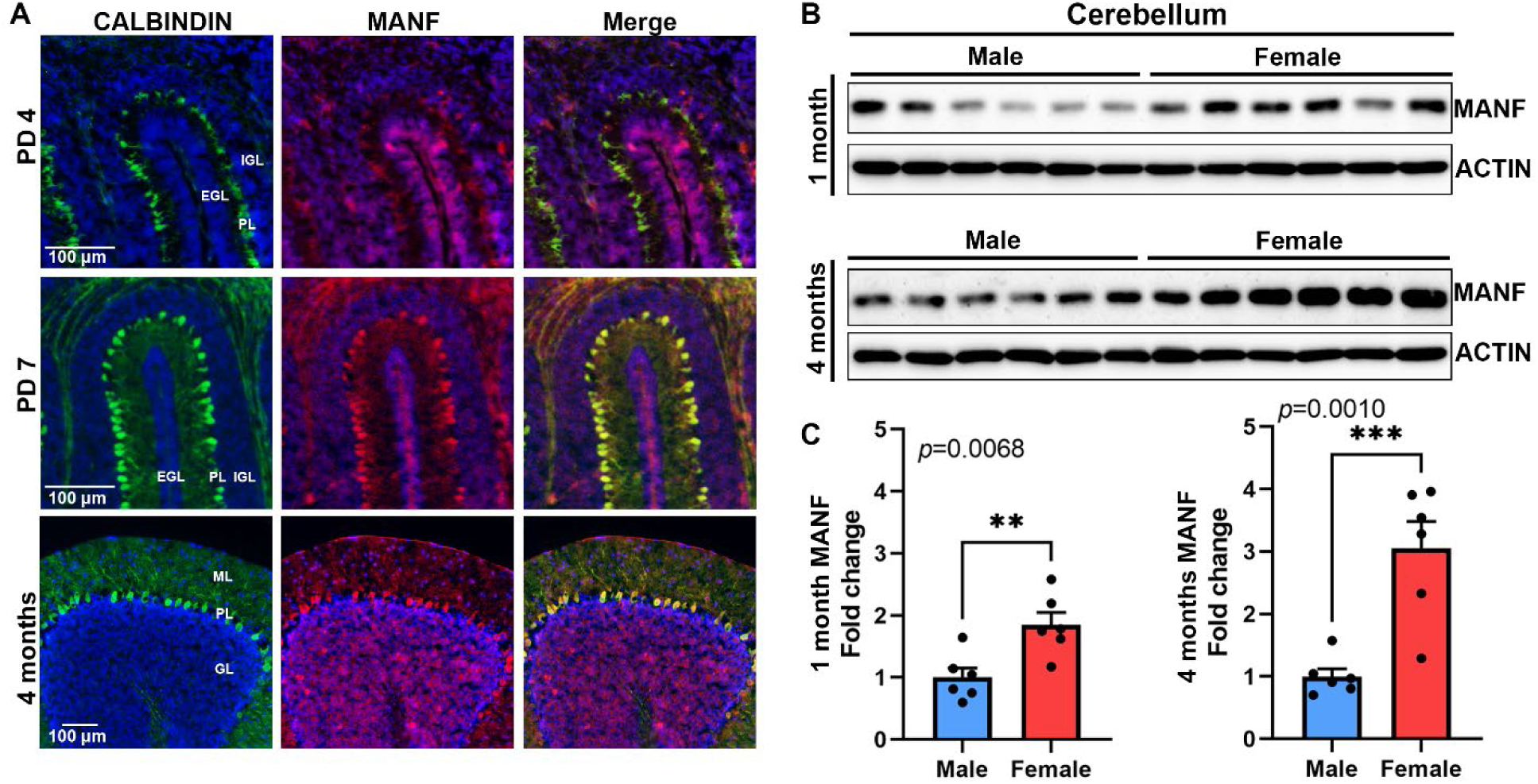
MANF expression in C57BL/6 wild type mice cerebellum. **A.** Representative immunofluorescent images of MANF (red) and PC marker Calbindin (green) in the C57BL/6 cerebellum at postnatal day (PD) 4, PD 7, and 4 months. Nuclei were counterstained with DAPI. EGL, external granular layer; IGL, internal granular layer; ML, molecular layer; PL, Purkinje cell layer; GL, granule cell layer. **B.** Representative immunoblot for MANF and ACTIN expression in 1- and 4-month-old C57BL/6 wild type male and female mice cerebellum. n=6 per group. **C.** Quantification of MANF expression normalized by ACTIN in 1- and 4-month C57BL/6 wild type male and female mice cerebellum. The data was expressed as mean ± SEM. n=6 per group. Student’s *t*-test. \*\**p*< 0.01; \*\*\**p*< 0.001.

**Figure S2.**
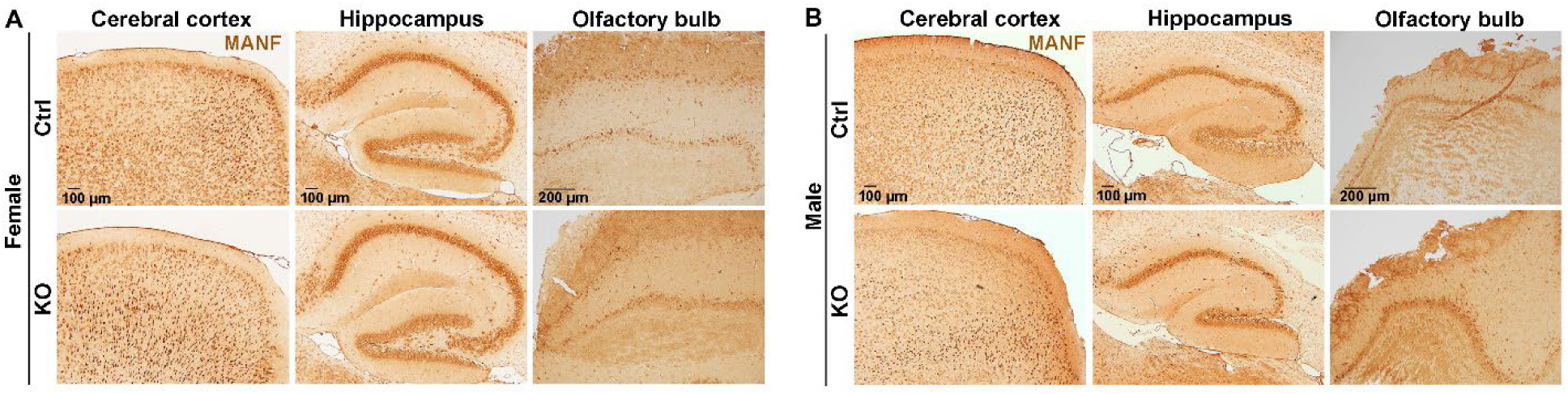
MANF expression in other brain regions is not affected. **A-B.** Representative immunohistochemistry images showing comparable MANF expression in the cerebral cortex, hippocampus, and olfactory bulb of adult female (A) and male (B) control and KO mice.

**Figure S3.**
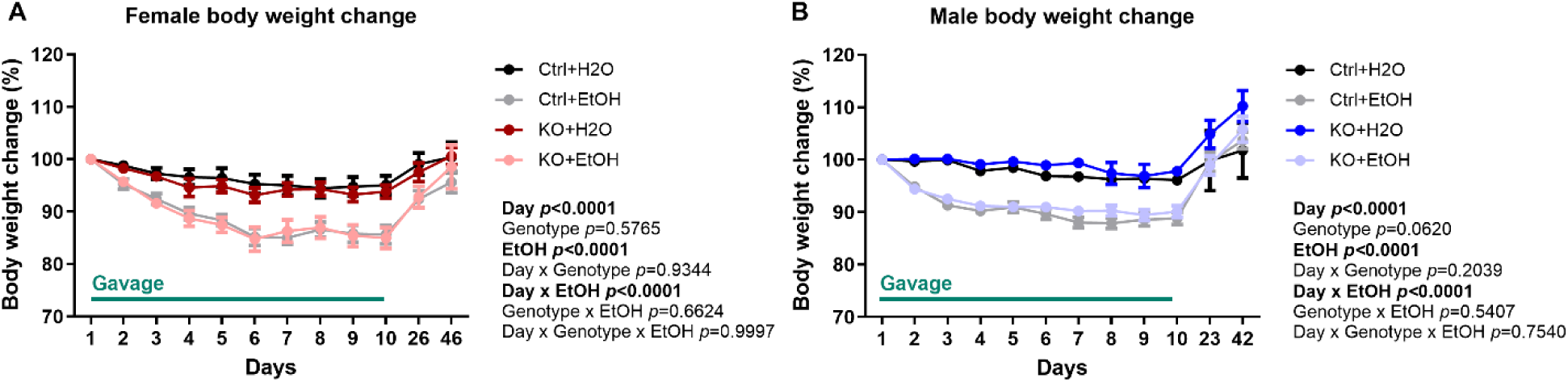
Mice body weight change during gavage and behavior tests. **A-B.** Body weight percentage change during gavage in female (A) and male (B). The data was expressed as mean ± SEM. n=8-11 per group. Three-way ANOVA followed by Tukey’s post hoc test. Significant main effect *p* values were highlighted in bold.

**Figure S4.**
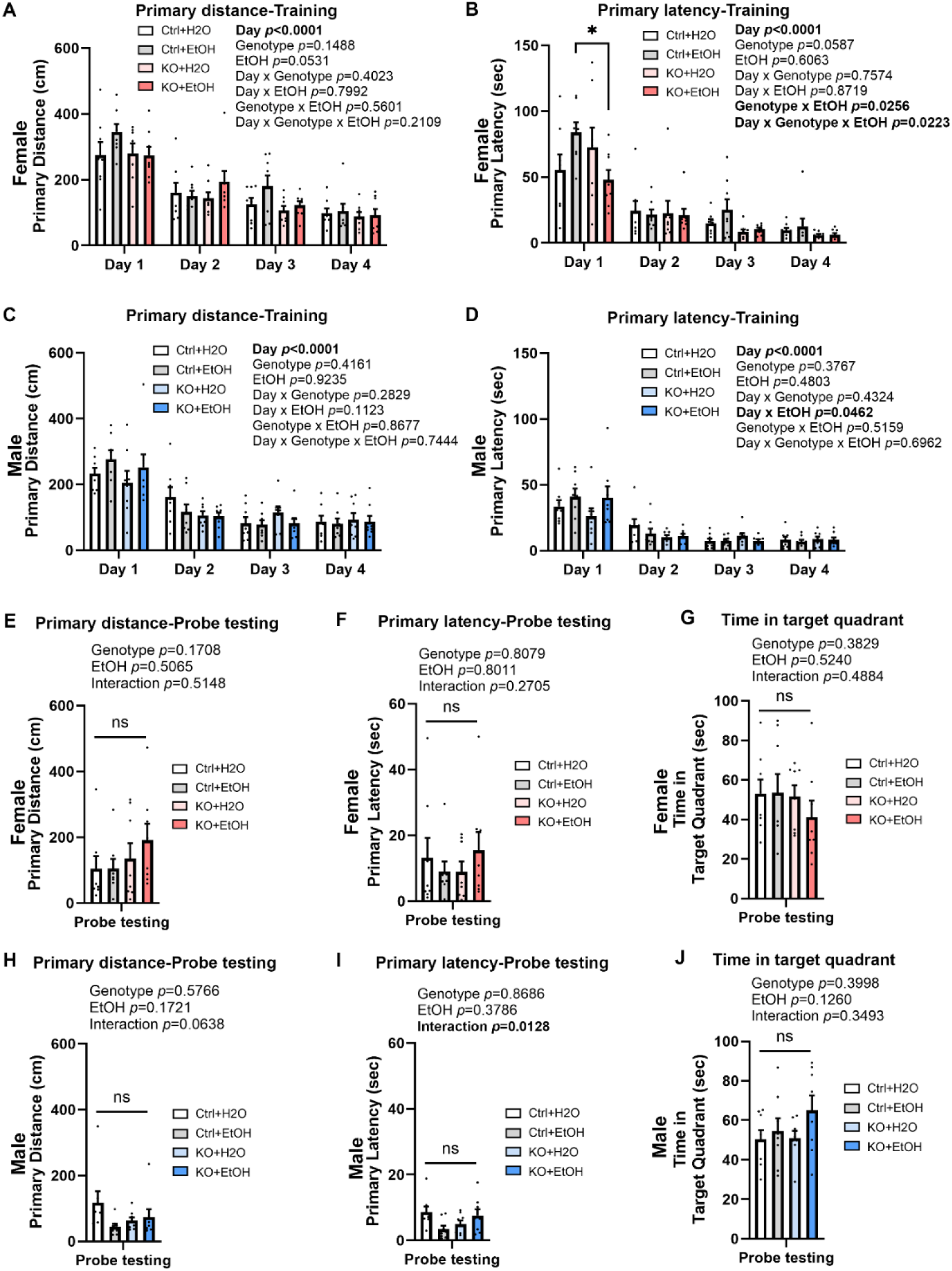
Effects of ethanol exposure on learning and memory in Barnes maze test. **A-D.** Primary distance traveled (A, C) and primary latency (B, D) to find the escape hole during training days in female (A, B) and male (C, D) mice. **E-J.** Primary distance traveled (E, H), primary latency (F, I) to find the escape hole, and time spent in the target quadrant (G, I) on probe testing day in female (E-G) and male (F-J) mice. All data were presented as mean ± SEM. n=8 per group. A–D, three-way ANOVA followed by Tukey’s post *hoc test*. E–J, two-way ANOVA followed by Tukey’s *post hoc* test. Significant main effect *p* values were highlighted in bold; \**p*< 0.05; ns not significant.

**Figure S5.**
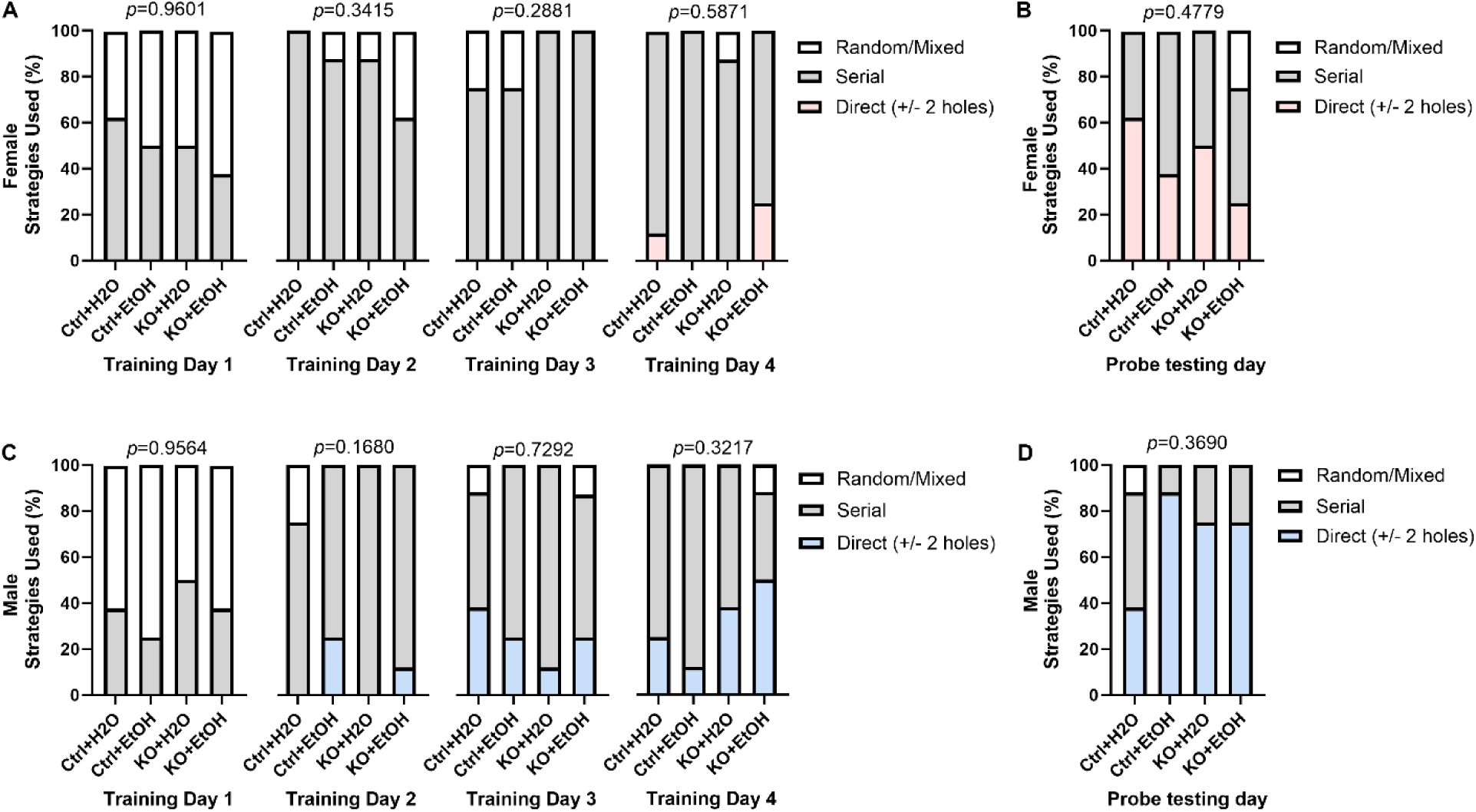
Searching strategies used by control and MANF KO mice in Barnes maze test. **A-B.** Percentage of each strategy used to find the escape hole during training (A) and probe testing period (B) in female. **C-D.** Percentage of each strategy used to find the escape hole during training (C) and probe testing period (D) in male. The three defined search strategies are random (top), serial (middle), and direct (bottom). n=8 per group. Fisher’s exact test.

**Figure S6.**
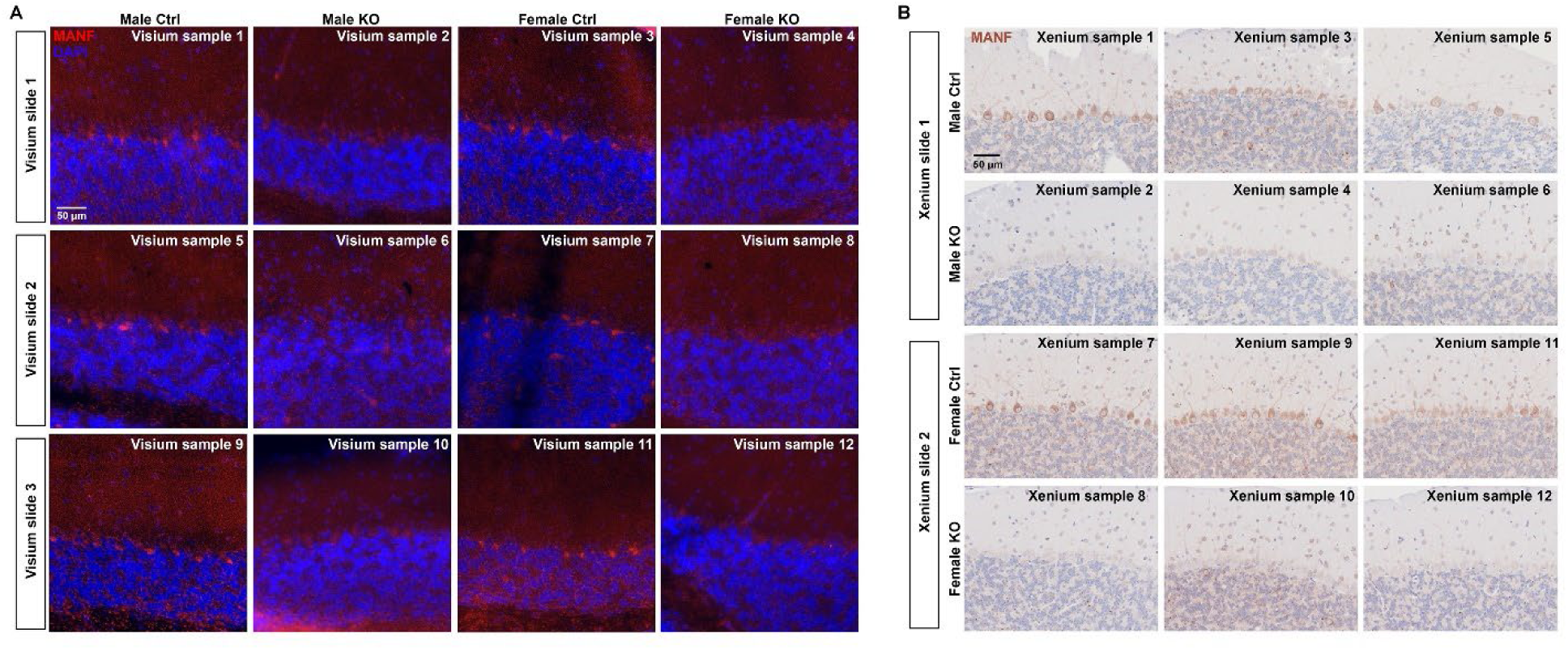
MANF expression in Visium and Xenium samples. **A.** Representative immunofluorescent images for MANF expression (red) in the 12 cerebellum samples used for Visium spatial transcriptomics analysis. Samples were counterstained with DAPI (blue). **B.** Representative immunohistochemistry images for MANF expression in the 12 cerebellum samples used for Xenium in situ analysis. Samples were counterstained with hematoxylin.

**Table S1.**
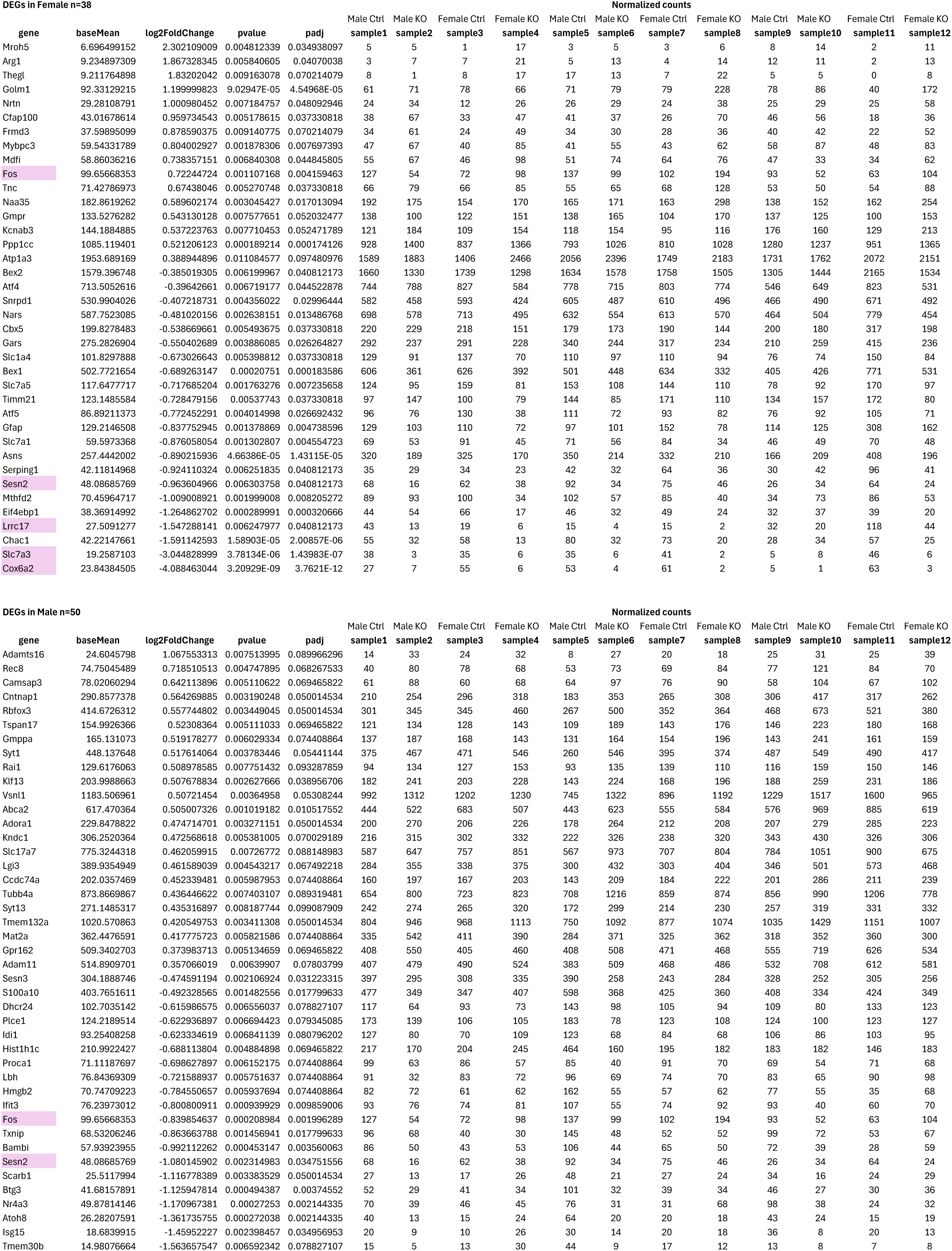
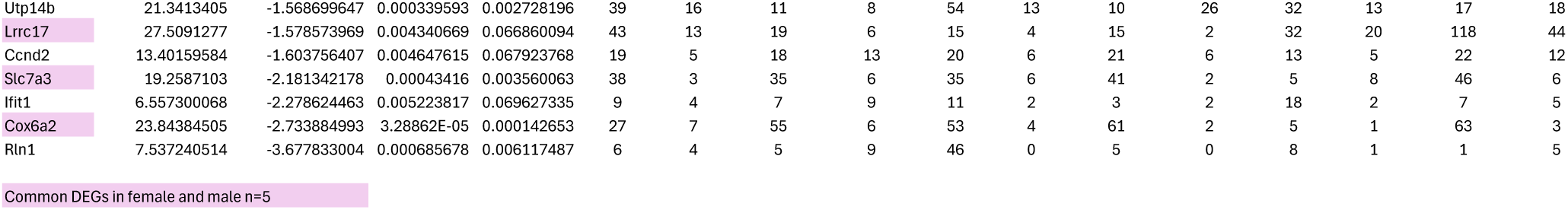
Visium PC cluster pseudobulk result.

**Table S2.**
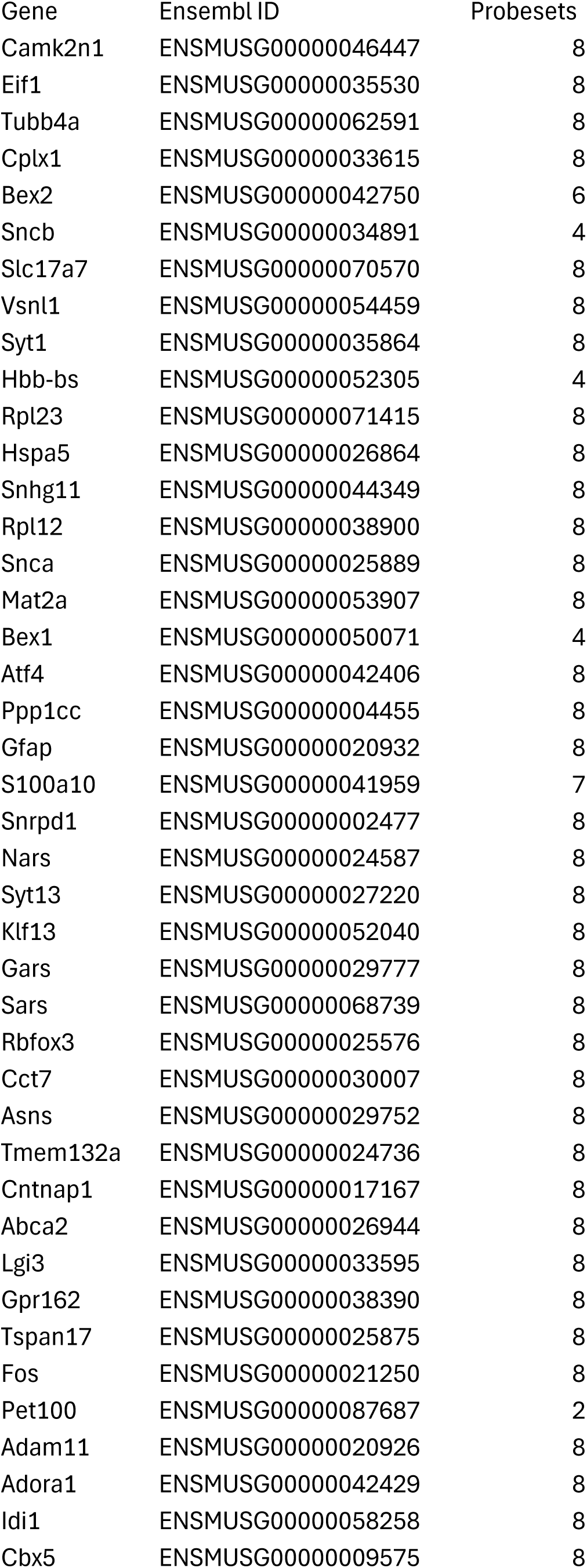
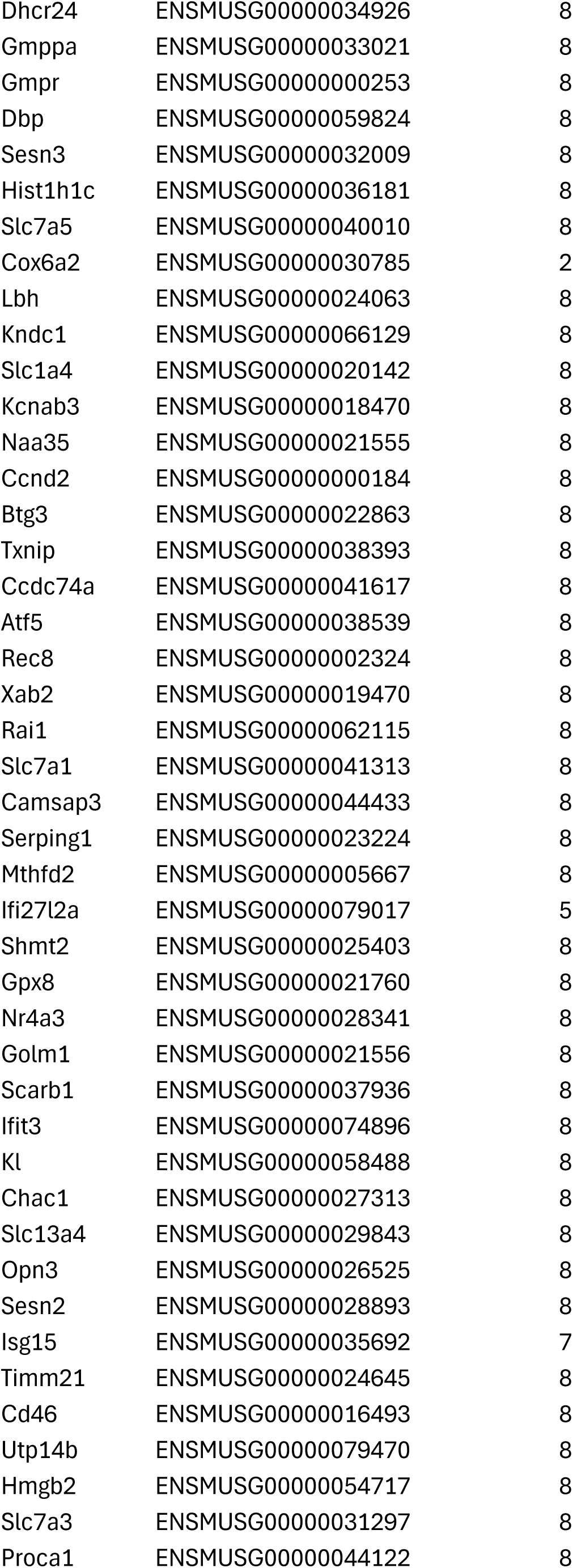
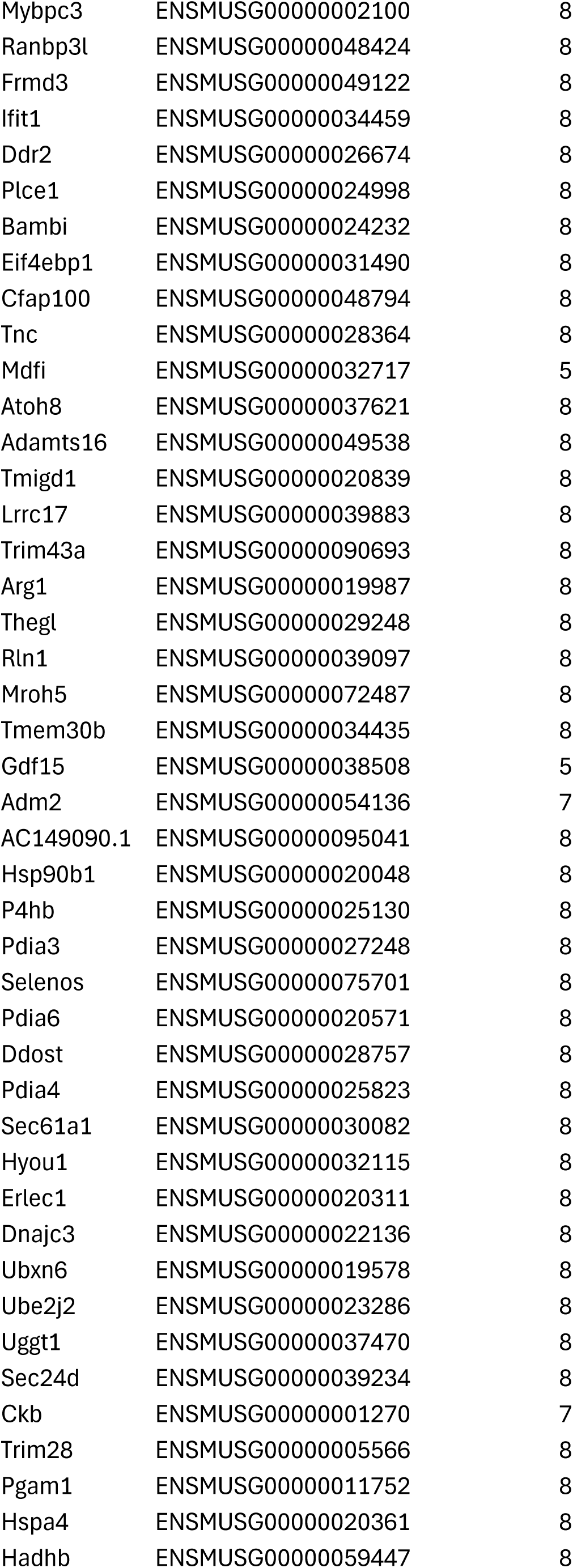
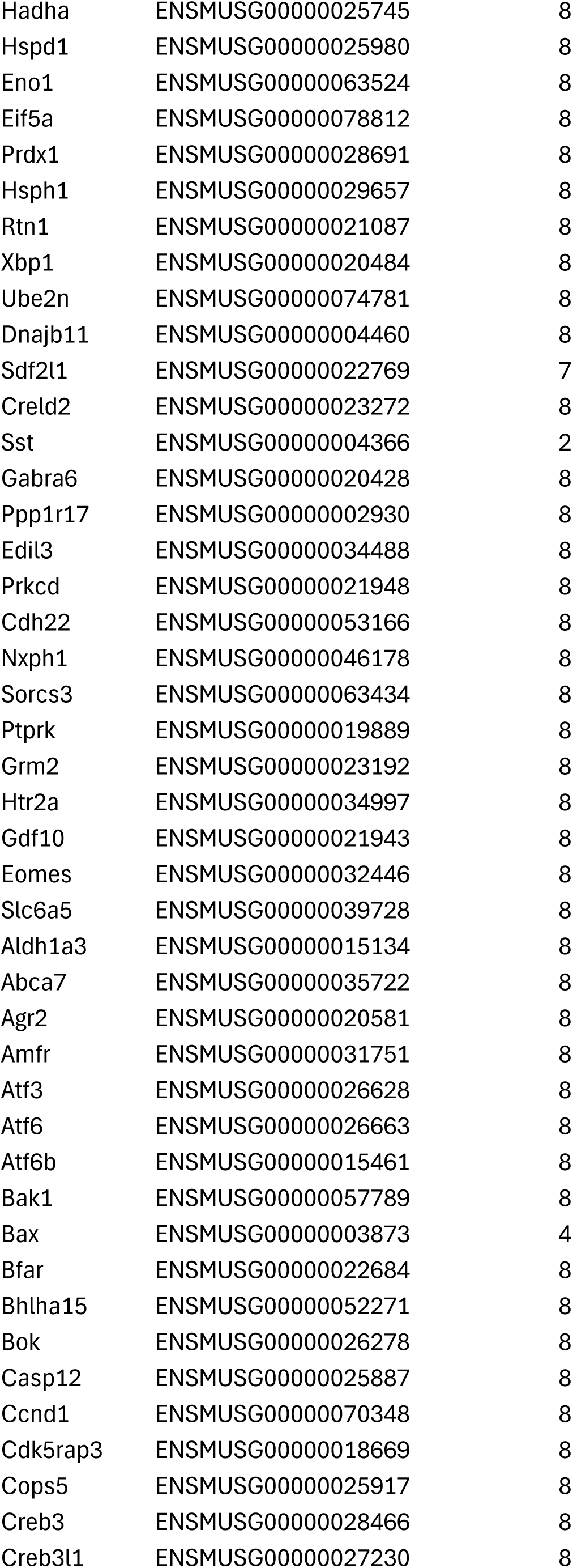
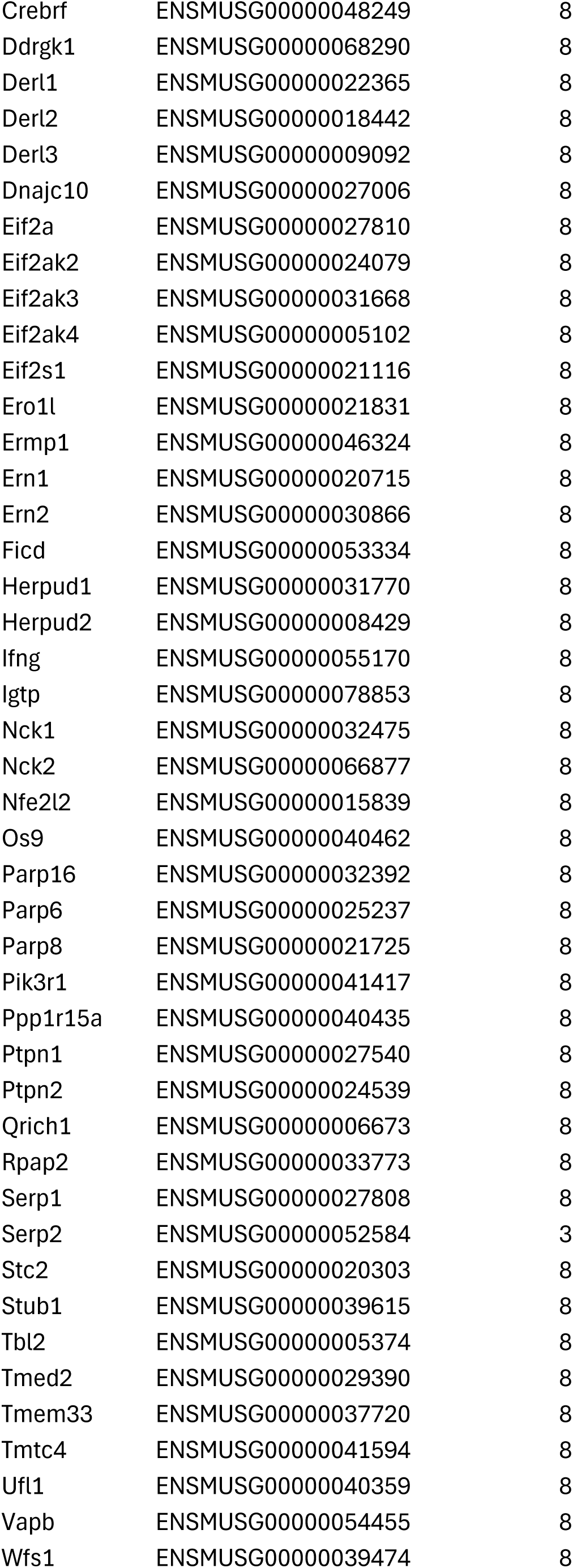
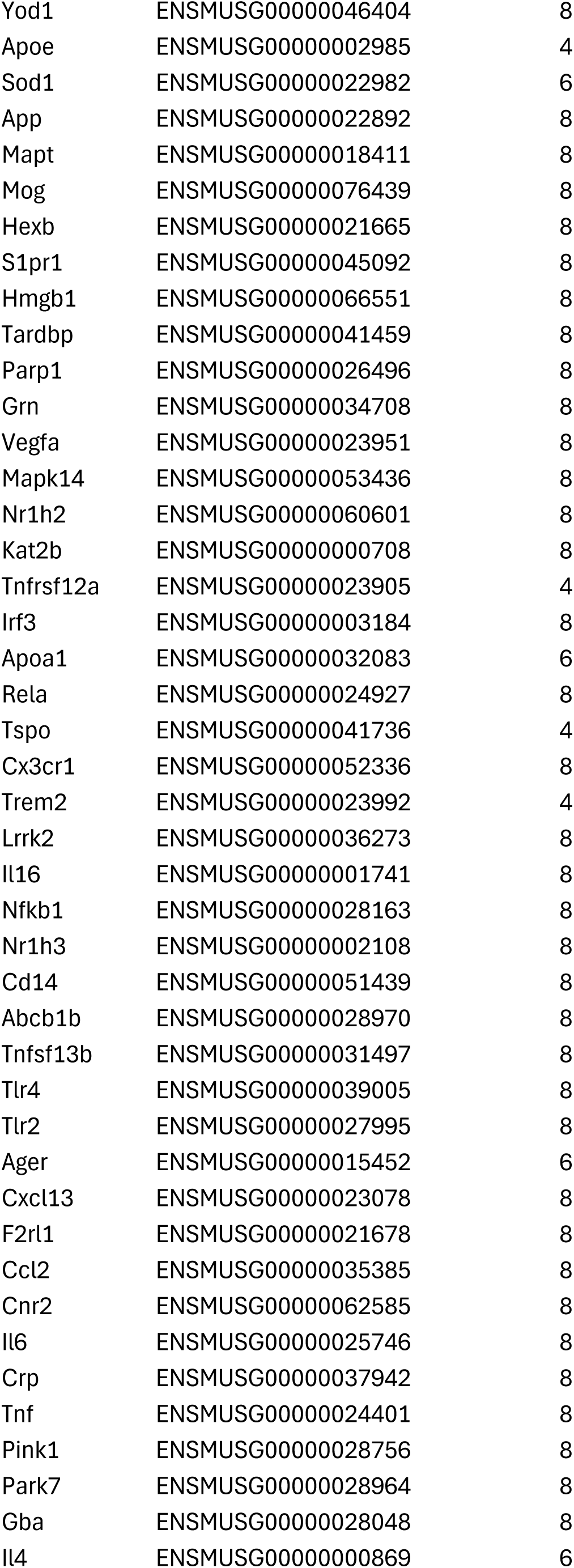
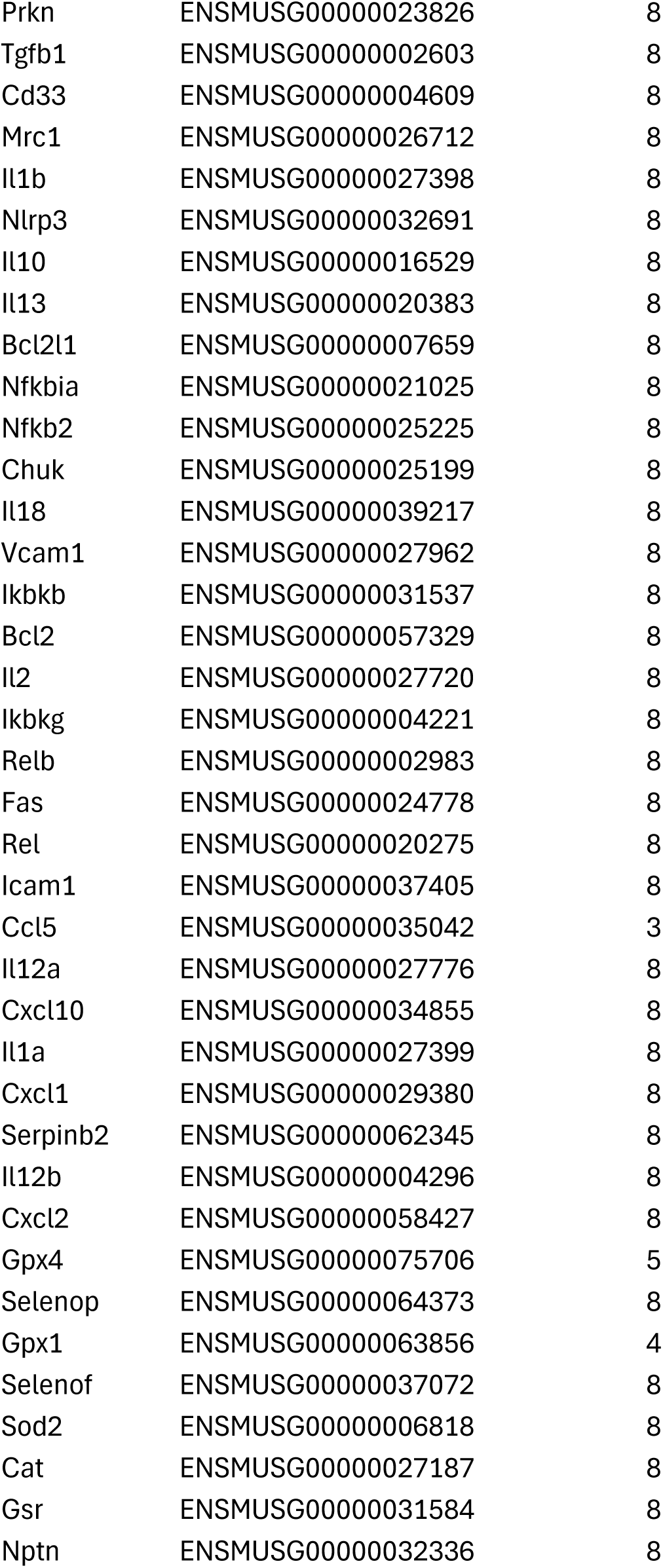
Xenium gene list.

**Table S3.**
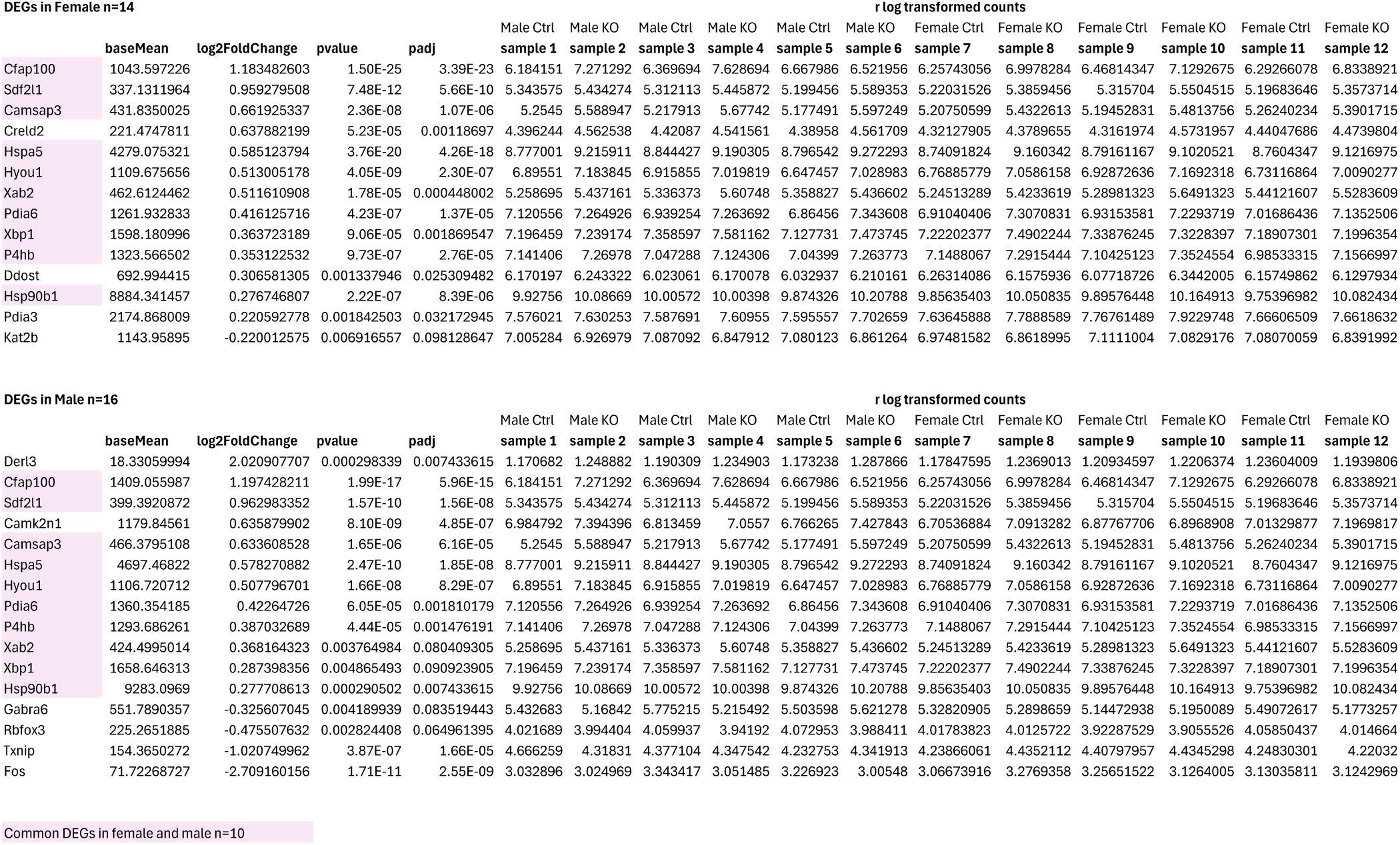
Xenium PC cluster pseudobulk result.

## Notes

### Competing Interest Statement

The authors have declared no competing interest.

